# Integration of Mechanistic Immunological Knowledge into a Machine Learning Pipeline Increases Predictive Power

**DOI:** 10.1101/2020.02.26.967232

**Authors:** Anthony Culos, Amy S. Tsai, Natalie Stanley, Martin Becker, Mohammad S. Ghaemi, David R. Mcilwain, Ramin Fallahzadeh, Athena Tanada, Huda Nassar, Edward Ganio, Laura Peterson, Xiaoyuan Han, ina Stelzer, Kazuo Ando, Dyani Gaudilliere, Thanaphong Phongpreecha, Ivana Marić, Alan L. Chang, Gary M. Shaw, David K. Stevenson, Sean Bendall, Kara L. Davis, Wendy Fantl, Garry P. Nolan, Trevor Hastie, Robert Tibshirani, Martin S. Angst, Brice Gaudilliere, Nima Aghaeepour

**Author notes:** Co-lead authorship (Anthony Culos, Amy S. Tsai). Co-senior authorship (Martin S. Angst, Brice Gaudilliere, Nima Aghaeepour).

## Abstract

The dense network of interconnected cellular signaling responses quantifiable in peripheral immune cells provide a wealth of actionable immunological insights. While high-throughput single-cell profiling techniques, including polychromatic flow and mass cytometry, have matured to a point that enables detailed immune profiling of patients in numerous clinical settings, limited cohort size together with the high dimensionality of data increases the possibility of false positive discoveries and model overfitting. We introduce a machine learning platform, the immunological Elastic-Net (iEN), which incorporates immunological knowledge directly into the predictive models. Importantly, the algorithm maintains the exploratory nature of the high-dimensional dataset, allowing for the inclusion of immune features with strong predictive power even if not consistent with prior knowledge. In three independent studies our method demonstrates improved predictive power for clinically-relevant outcomes from mass cytometry data generated from whole blood, as well as a large simulated dataset.

## 1. Introduction

In response to an immunological challenge, immune cells act in concert to form a complex and dense cell-signaling network^1,2^. The single-cell evaluation of intracellular signaling responses is particularly valuable in characterizing this cellular network as it provides a functional assessment of an individual’s immune system. In clinical settings, a deep understanding of functional immune responses not only provides diagnostic opportunities, but also is often the first step in developing immune therapies (recent examples include successful immune modulation in chronic lymphocytic leukemia^3^, neurodegeneration^4^, and Ebola^5^).

Advanced flow cytometry technologies can characterize millions of single cells from a given patient, which enables the identification of signaling pathways even in rare cell populations^6^. The recent advent of high-dimensional polychromatic flow cytometry^7,8^ and mass cytometry^9,10^ technologies have vastly increased our ability to study the human immune system with unprecedented functional depth by increasing the number of features measured per cell. However, the increased dimensionality, small cohort sizes in clinical studies, and the inherently complex networks of internal correlations between the measured cell-types and pathways presents unique computational challenges^11^. Translating these immunological observations into clinically relevant mechanisms requires statistically rigorous analysis techniques. Multivariate modeling, in contrast to univariate analysis, can simultaneously consider all measured aspects of the immune system to increase predictive power. However, multivariate modeling requires exponentially larger cohort sizes as the number of measurements grow (*a.k.a.*, “Curse of Dimensionality”^12–14^); This is especially true for more powerful deep-learning based models which provide greater predictive power but require substantially larger sample sizes. In practice, increasing the cohort size by several orders of magnitude to power such analyses is often a significant challenge in clinical settings. Moreover, multivariate analyses performed on all available measurements produce large complex models that are difficult to implement in resource constrained settings^15^ and often lack robustness^16^.

Integration of prior knowledge has been broadly recognized as an effective approach for reducing model complexity and increasing robustness^17–21^. In biological sciences, examples of such knowledge integration include inference of biological networks^22^ and causal pathway modeling^23^. In modern immunological datasets, however, integration of prior knowledge has been impractical due to the unstructured format of prior immunological datasets and the complex nature of the measured features. In this work, we propose a framework for integration of prior immunological knowledge into the model optimization process of the Elastic Net^24^ (EN) algorithm (Fig. 1). In our immunological Elastic-Net (iEN) framework, the prior knowledge developed by a panel of expert immunologists on a per-feature basis is integrated into the EN algorithm as feature weights during coefficient optimization (similar to adjusting Bayesian priors - see the Methods section). The addition of knowledge-based immunological priors guides the sparsification process to solutions more consistent with biological knowledge while still allowing all measured immune features to be included in the exploratory analysis. In our experiments, the iEN outperformed the standard EN, as well as a broad range of standard machine learning algorithms. A two-layer repeated 10-fold cross-validation (CV) was used to determine model consistency and establish a clear comparison of model performance between the algorithms. With the first CV layer the free parameters of the models are optimized (including a factoring controlling the impact of domain knowledge in the case of the iEN) and the second CV layer predicted previously unseen observations.

**Fig. 1.**
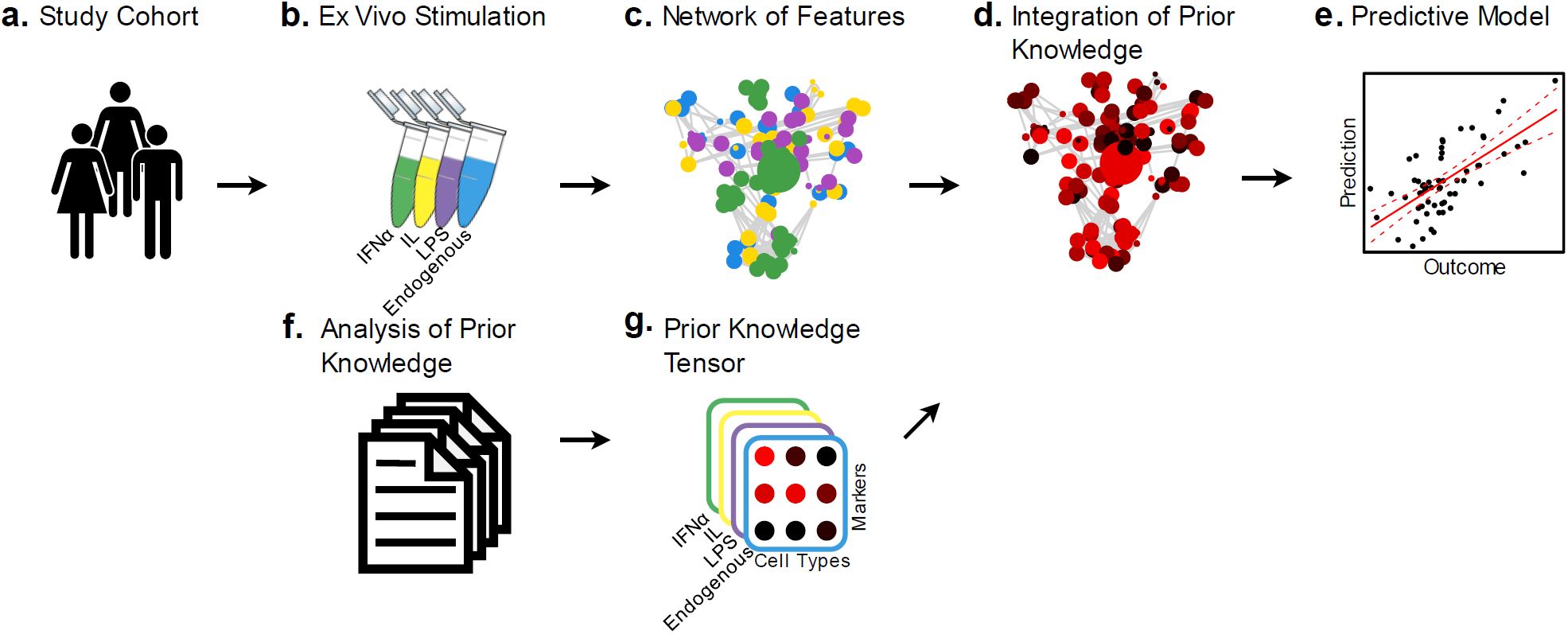
The immunological Elastic-Net analysis pipeline: **(a)** Individuals within the cohort of study provide blood samples, which is subsequently **(b)** stimulated with ligands *ev vivo* to activate various functions of the immune system. **(c)** This produces single cell measurements of the immune system, resulting in a complex network of cell types and signaling pathways representing both innate and adaptive immunity. **(d)** This dataset is then fed into the iEN algorithm for **(e)** predictive modeling of the outcome of interest. **(f)** Immunological prior knowledge for each feature, in response to each *ex vivo* stimulation condition is extracted by a panel of experts and encoded into a prior knowledge **(g)** tensor to guide the model optimization process.

In this article, we have included two real-world clinical examples as well as a large simulation study: The first analysis, as an example of a continuous clinical outcome, identified components of maternal immune adaptations in a Longitudinal Term Pregnancy (LTP) study, which included a blinded validation cohort. The second example was a classification analysis of a categorical outcome, modeling patient and control populations for Chronic Periodontitis (ChP). The third example used synthetic data generated to replicate mass cytometry measurements to enable in-depth understanding of the iEN behavior across varying cohort sizes. Additional analyses were run to determine the effect of prioritization on general model behavior and the stability of results given errors within the prior knowledge in simulated and real world data. Each of the three examples were chosen to determine the generalizability of the iEN algorithm, as well as its efficacy, in a range of real-world scenarios.

## 2. Results

### 2.1. Integration of Prior Immunological Knowledge into a Multivariate Model

The biological priors used in the iEN model were created by an independent panel of immunologists, such that features more consistent with known biology have higher values (Supplemental Tables 1 and 2). Prior knowledge tables constructed before the analysis emphasized receptor-specific signaling responses describing canonical pathways activated downstream of *ex vivo* stimulation conditions used in the mass cytometry assays. For example, panel members broadly agreed on the prioritization of the phosphorylation of STAT1, STAT3 and STAT5 in all adaptive and innate immune cells in response to IFNα stimulation^25,26^; the phosphorylation of STAT1, STAT3, STAT5, and ERK1/2 MAPK in all adaptive and innate immune cells in response to the IL cocktail containing IL-2 and IL-6^27,28^; and the phosphorylation of P38 MAPK, MAPKAPK2, ERK1/2, rpS6, CREB, and NF-κB and total IκB signal in all innate immune cells (except pDCs), and in regulatory T cells in response to LPS stimulation condition^29–33^. An example of all measured immune features and those selected by this prior knowledge tensor is presented in Fig. 2.

**Fig. 2.**
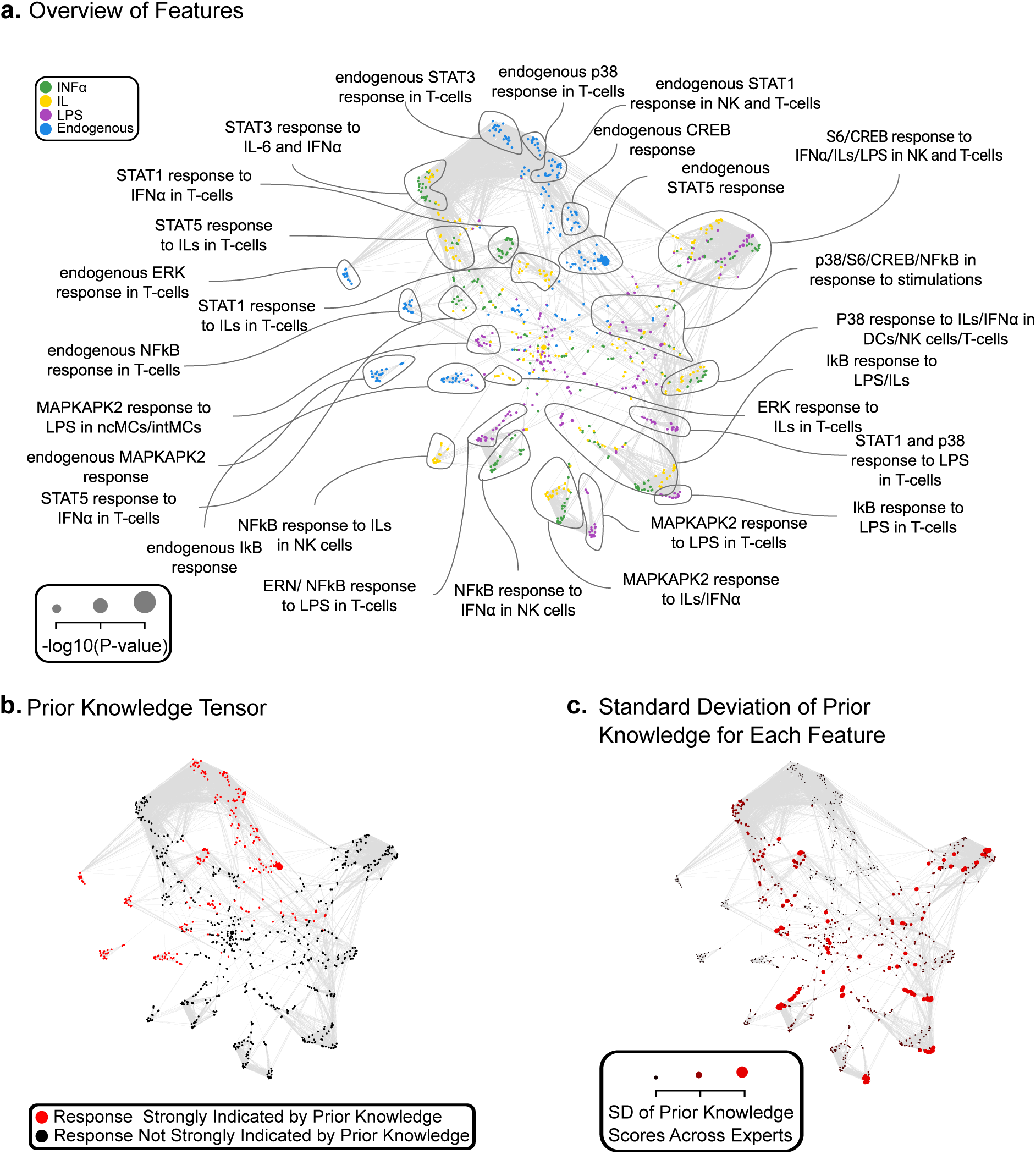
Integration of immunological priors: Overview of the the LTP study. **(a)** A correlation network of intracellular signaling responses, measured in peripheral immune cells, colored by *ex vivo* stimulation status. Edges represent significant (P-value < 0.05) pairwise correlation after Bonferroni correction. Node sizes represent the significance of correlations with the response variable (gestational age of normal pregnancy). **(b)** Immune features that were congruent with domain specific knowledge as determined by a panel of five immunologists are refined into a tensor and then projected onto the correlation network. Here, immune features which have a value of 1 (full agreement among the panel) are colored red while all other immune features are colored black. **(c)** The network is colored by the standard deviation of scores assigned to each feature by the panel of immunologists. The consistency of the panel scores was generally higher amongst the features with a higher score, indicating more diversity in the scores assigned by the panel for exclusion of features and a stronger agreement regarding the top features that should be prioritized by the algorithm.

These scores vary from zero to one, with one representing the immune features that are most consistent with prior knowledge according to the panel of experts. iEN regularized regression models are constructed through optimization of the objective function *L*(*β*) = ‖*Y* − *Xϕβ*‖^2^ + *λ*[(1 − *α*)‖*β*‖^2^/2 + *α*‖*β*‖_1_] where *X* is a matrix of *p* measured immune features (columns) for *n* patients (rows) and *Y*is a vector of the clinical outcomes of interest. The algorithm calculates the coefficients *β* which minimize the objective function subject to the *L*_1_ = ‖*β*‖_1_ and *L*_2_ = ‖*β*‖^2^ penalties. The combination of these penalty terms allow for the selection of the features correlated with the outcome of interest and the exclusion of redundant measurements, while also accounting for internally correlated measurements. Model optimization for iEN is controlled by three parameters, *λ, α*, and *φ*. These parameters can be interpreted as the amount of sparsity in a model (*λ*), how sparsity is balanced between the *L*_1_ and *L*_2_ penalty terms (*α*), and the amount of prior knowledge prioritization (*φ*). Prioritization of biologically consistent features is accomplished through *ϕ*, which is a *p* × *p* diagonal matrix of the form *diag*(*ϕ*) = {*φ*_1,1_, *φ*_2,2_, …, *φ*_*p,p*_}such that 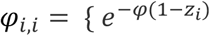 where *z*_*i*_ is the score of the *i*^*th*^ immune feature, and *φ* is the amount of prioritization attributed to the model}. This definition allows for a limited effect of *φ* on model coefficients while also increasing the impact of the features consistent with the prior knowledge tensor (Fig. 3). Similar to the *α* and *λ* free parameters, prioritization of prior knowledge affects the sparsification (Fig. 3 & Supplemental Fig. 1) and optimization of the model (Supplemental Fig. 2). The two-layer 10-fold CV used for iEN optimization and estimation was implemented and parallelized over the parameters *α, λ*, and *φ*. Runtime analysis for this procedure is presented in Supplemental Fig. 3b.

**Fig. 3.**
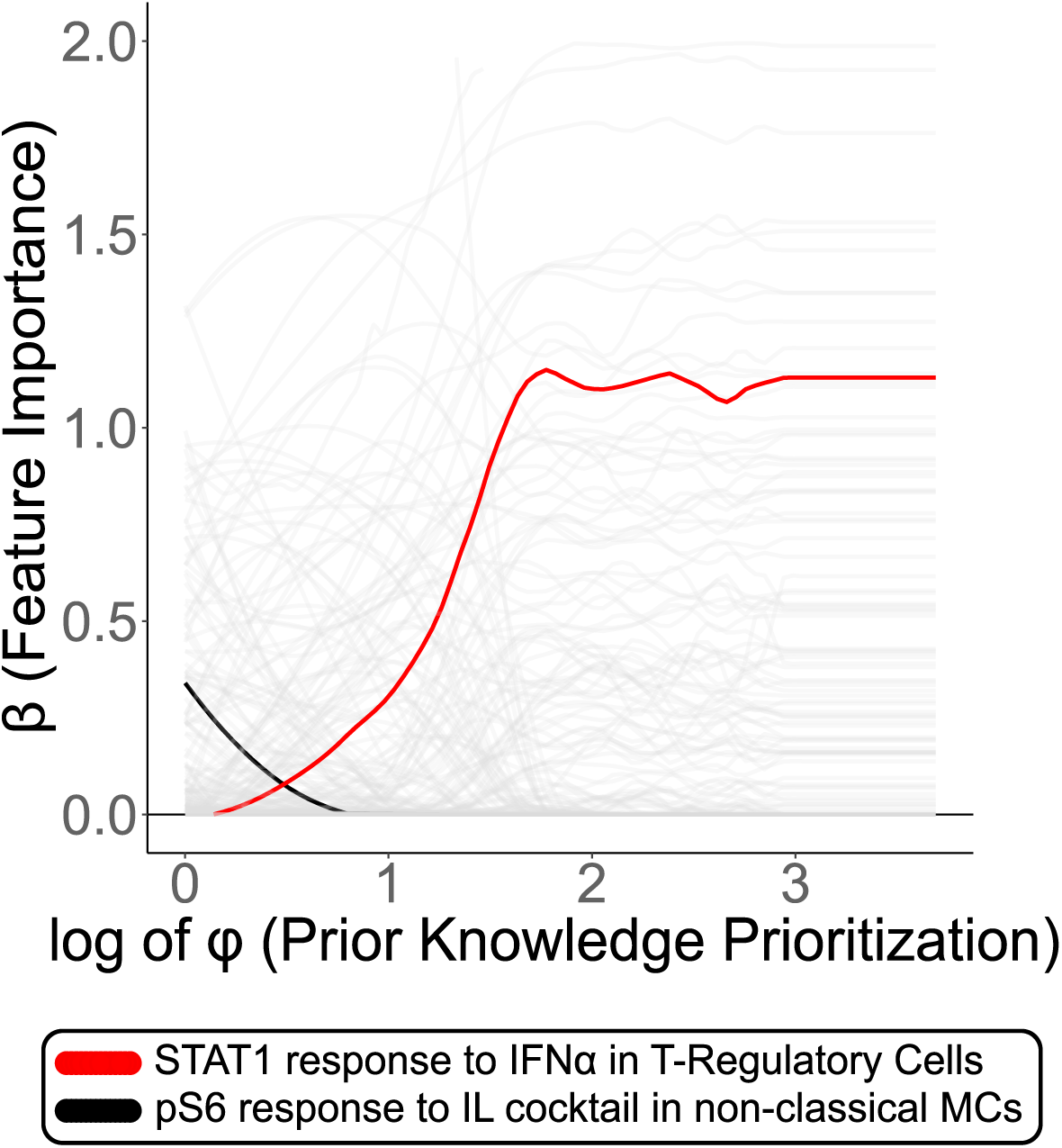
Prior scores effect on sparsification: An example of the impact of prior immunological knowledge on various features in an iEN model is visualized. As *φ* is increased across the X-axis (increased impact of prior knowledge), the contributions of each feature to the final model (y-axis) changes to select models consistent with immunological priors. Two examples are highlighted where a feature is emphasized or de-emphasized (in red and black, respectively) by prior knowledge. In this example, the STAT1 response to IFNα stimulation in regulatory T-cells is prioritized as STAT1 downstream of the IFN-α/β receptor is integral for their homeostasis and function34. Conversely, the prpS6 response to stimulation by IL-2 and IL-6 in ncMCs is progressively deprioritized as this signaling response is inconsistent with prior understanding of these signal transduction pathways in this cell-type; IL-2 primarily drives T-cell differentiation through the JAK/STAT pathway^35^. Similarly, IL-6 primarily activates the JAK/STAT pathway and IL-6 receptors are expressed only in a subset of immune cells^36,37^. This confirms that integration of the priors not only can modify the algorithm’s behavior, but also that the intensity of this impact can be controlled through the *φ* free parameter.

### 2.2. Example 1 - Analysis of Gestational Age during Longitudinal Term Pregnancy

The first example investigated the adaptations of the maternal immune system during pregnancy that can be incorporated into predictive models of gestational age^38^. During a healthy pregnancy, the immune system strikes a delicate balance to enable tolerance towards the fetus and simultaneously mount a response to defend against pathogens. Abnormal immune system adaptations during pregnancy have been linked to adverse maternal and neonatal outcomes, such as pregnancy loss, preterm birth, and preeclampsia^39–41^. This study aims to understand the immunological mechanisms behind term birth as a pivotal first step in understanding abnormal pregnancies and their impact on long-term outcomes ^42^. In this example, a total of 54 blood samples from 18 women were studied during and six weeks after pregnancy. Three antepartum blood draws were collected at different gestational ages, with gestation being measured via ultrasound at time of sample collection. The resulting 54 whole blood profiles of the immune system were manually gated (Supplemental Fig. 4) into 24 cell types to measure endogenous activity of 10 signaling markers, as well as activity of these 10 markers in response to *ex vivo* stimulations with three different ligands, providing 960 immune features for analysis (Fig. 2a). Prediction of gestational age in this dataset was a significant challenge as evident by t-Distributed Stochastic Neighbor Embedding (t-SNE)^43^ analysis which displayed no easily identifiable patterns in a two dimensional projection (Supplemental Fig. 5a). Predictive models were built using iEN and model parameters were optimized to minimize the residual sum of squares of the predicted versus actual gestational age. iEN produced models of immune features that more accurately predicted gestational age than the similar EN analysis, and other contemporary machine learning methods, which were agnostic to the immunologic priors (Fig. 4a, 4e & Supplemental Fig. 5b, and 6a). iEN analyses of postpartum samples demonstrated that the immune system returns to a state similar to early pregnancy by six weeks postpartum (Supplemental Fig. 5c). An additional cohort of ten women were prospectively studied and analyzed as a blinded validation set. Importantly, the blinded validation cohort also demonstrated that iEN models produced substantially more accurate results than the EN algorithm (Fig. 4b, 4f & Supplemental Fig. 5e, and 5f). Stepwise reduction of iEN and EN model coefficients revealed superior predictions in the validation cohort by the iEN compared to the EN algorithm for models of equal size (Supplemental Fig. 5d).

**Fig. 4.**
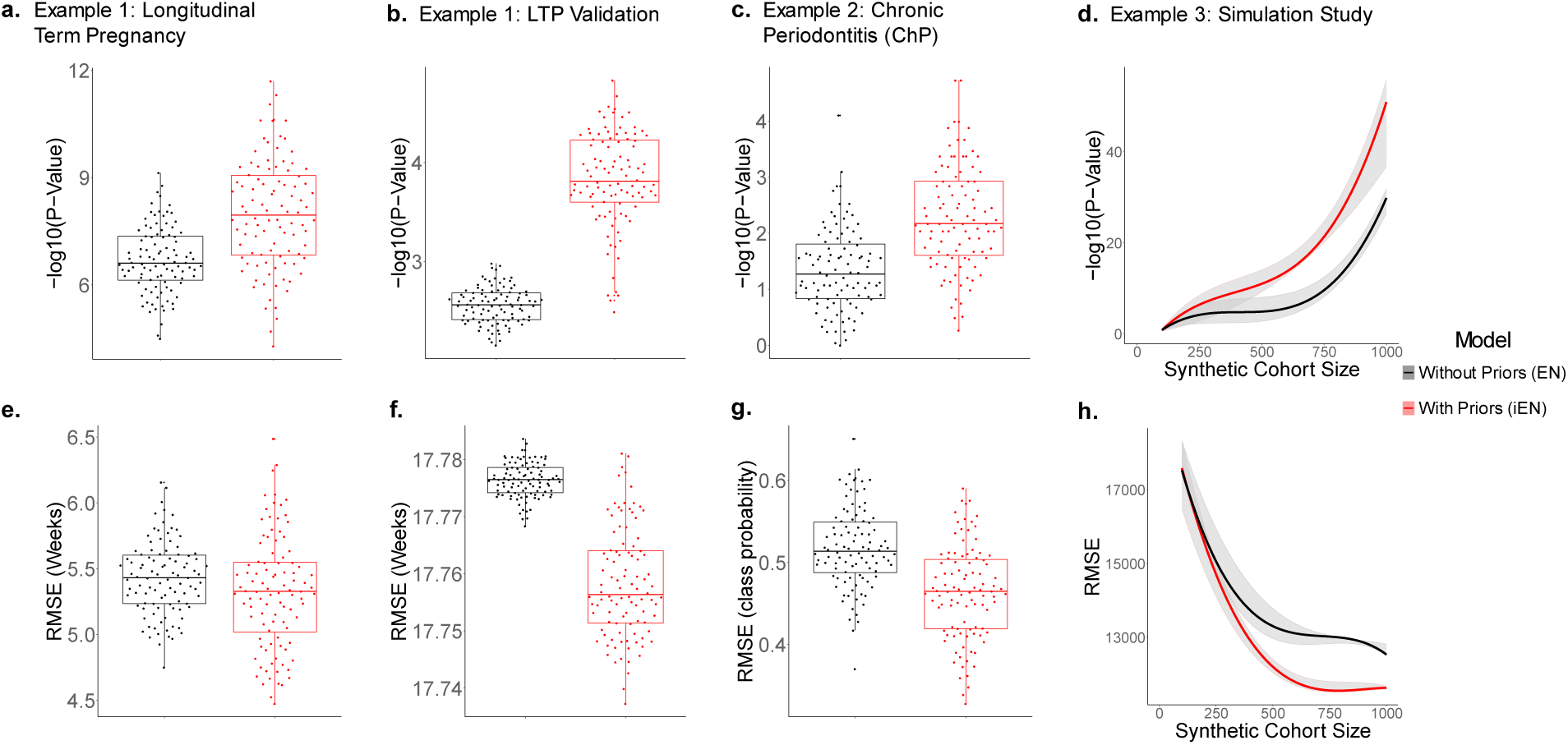
Incorporation of prior knowledge improves predictive power in two clinical studies and a simulated experiment: **(a)** Boxplot of Pearson correlation P-values calculated on out-of-sample predictions from repeated 10-fold cross-validation of EN (black) and iEN (red) models for the LTP dataset. **(b)** Validation of the LTP model on an independent validation cohort. These predictions are compared against the true response variable via -log10 Pearson correlation P-value. **(c)** Boxplots of Wilcoxon Rank-Sum test P-values similarly calculated on out-of-sample predictions for the ChP dataset. Comparison of EN and iEN model performance for the respective datasets demonstrated improved predictive power for the iEN exhibited by -log10 P-values. **(d)** Simulated study with varying cohort sizes of simulated “patients” with 700 features demonstrated a larger gain (measured by -log10 Pearson’s test P-value) for integration of prior immunological knowledge in datasets with a relatively small cohort size and a large number of features. Median predicted value over multiple cohort sizes is displayed here with the interquartile range shaded in gray. To demonstrate the effect size of these models, RMSE values are visualized in panels **(e), (f), (g)**, and **(h)**, respectively.

### 2.3. Example 2 - Analysis of Chronic Periodontitis

The second example investigated the classification of patients with ChP, a chronic inflammatory disease of the oral cavity. ChP is associated with severe systemic illnesses (such as heart disease, various malignancies, and preterm labor)^44,45^ and, in its most severe form, affects approximately 11 percent of the global population^46^. A better understanding of the immunological manifestations of ChP is a critical first-step for the development of immune therapies that may alter the course of systemic diseases associated with ChP. This dataset was generated from 28 participants, 14 diagnosed with ChP and 14 healthy controls. Blood samples from the study participants were analyzed by mass cytometry and manually gated (Supplemental Fig. 4) for 18 cell types to measure 11 signaling markers in response to *ex vivo* stimulation with 4 different ligands; this provided a total of 792 immune features for analysis (Supplemental Fig. 7a). Application of t-SNE demonstrated that the two study populations (patients and controls) were not easily separable (Supplemental Fig. 7b), thus motivating the use of supervised predictive modeling. For this example, iEN and EN algorithms were used to fit binomial models for classification of patients and controls. All free parameters were optimized for the Area Under the Receiver Operator Curve (AUROC)^47^. Analysis results indicated that iEN outperformed the EN (Fig. 4c, 4g & Supplemental Fig. 7c) as well as other machine learning methods (Supplemental Fig. 6c).

### 2.4. Example 3 - Simulation Study

The third example, a simulation study, demonstrated the particular advantages of prior knowledge in studies with limited cohort sizes and numerous features. Datasets were generated with 700 features per simulated patient while the number of patients varied from 100 to 1000 in increments of 100. Data were generated with features that are random and uniform. A limited number of features were generated to have various degrees of correlation with the response variable. Specifically, 50 highly predictive features, 200 moderately predictive features, and 450 randomly distributed features were assigned a corresponding biological prior. For a more detailed description of the data generating process see the simulated data section in Methods. Repeated 10-fold analysis of the simulated data with increasing population sizes displayed a convergent trend between the iEN and EN models as *n* increased (Fig. 4d, 4h, & Supplemental Fig. 3a). These results indicated that integration of prior knowledge is of particular importance in clinical systems immunology settings where the cohort size is limited yet a relatively large number of immunological features are measured.

### 2.5. Sensitivity Analysis of the Prior Knowledge Tensor

The iEN pipeline depends on the prior knowledge tensor. We therefore investigated the iEN’s robustness by introducing errors into the prior knowledge tensor of each of the three examples. Introduction of moderate to substantial error into the prior tensor resulted in a consistent reduction in the predictive benefit for the iEN over the traditional EN model. Error was introduced stochastically and progressed towards uniform random noise, with 11 total incremental steps and 100 tensors generated per increment. As the prior tensor approaches a uniform random distribution (the highest amount of error in the prior knowledge matrix) the iEN and EN performances converged. These results remain consistent across the LTP, LTP validation, ChP, and simulation studies (Fig. 5). This robustness against induced error demonstrated across multiple scenarios displayed the practicality of the iEN, given that some amount of error in the prior knowledge is to be expected in real world applications. Such robustness allows for moderate amounts of error and disagreement between experts when quantifying the biological consistency of features.

**Fig. 5.**
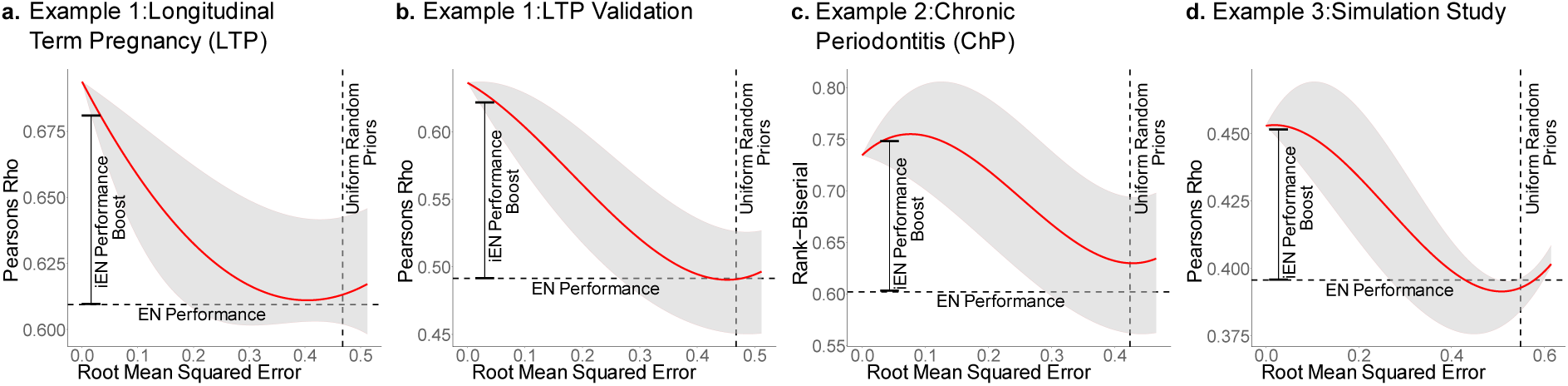
iEN is robust to errors in the prior knowledge tensor: Various levels of noise were artificially added to the prior knowledge values, as indicated by the Root Mean Squared Error of the true prior values vs. the simulated ones (x-axis). As the value on the x-axis increases, the amount of noise in the simulated prior increases until all priors are sampled from a random uniform distribution (vertical dashed line). Reassuringly, at this point iEN’s performance is close to EN’s performance (with no priors) as indicated by a horizontal dashed line. Importantly, iEN continues to outperform EN (indicated by the horizontal dashed line) for even high amounts of error in the priors.

### 2.6. Empirical Evaluation

In all analyses, the integration of expert knowledge improved the prediction of clinical outcomes in comparison against EN with no prior knowledge (Fig. 4) as well as standard machine learning algorithms including EN ^24^, LASSO^48^, RF^49^, SVM^50,51^, and KNN^52^ (Supplemental Fig. 6). Further comparison of iEN and EN models by features selected demonstrated substantial overlap between the models; however, model comparison by coefficient weights displayed a substantial difference in predictive importance of the features selected (Supplemental Fig. 8). Hyper parameter selection frequency across all models generated displayed a consistent prioritization of prior knowledge via *φ* (Supplemental Fig. 9). The integration of prior knowledge allows the iEN to determine the predictive benefit of prioritizing canonical signaling pathways in a data driven manner (Supplemental Fig. 2). Importantly, this enables iEN to function without excluding any of the features from consideration even when scored as inconsistent with prior knowledge by the human experts. This behaviour allows for the iEN to contain the EN as an edge case when the prior knowledge is not beneficial. Similarly when prioritization is most beneficial the resulting model will contain only features scored as 1 in the prior tensor (those with complete consensus among the panel of experts).

## 3. Discussion

A structured collaboration between clinicians, biologists, and computer scientists can lead to machine learning algorithms in life sciences achieving stronger results ^53^. In this article, we proposed a collaborative framework that enables integration of prior knowledge of cell signaling pathways in a machine learning algorithm to increase the predictive power and robustness of the resulting models in clinical datasets. The iEN improved accuracy of clinical predictions in multiple scenarios, even when moderate amounts of noise were artificially added to the extracted prior knowledge data. These benefits are especially evident in settings with small cohort size and a large number of measured features, as is common in modern systems-level clinical studies. The data-driven approach implemented allows for prior knowledge only to be incorporated when a predictive benefit is observed. Functionally, this reduces the regularization of the features consistent with prior knowledge, resulting in the development of sparse models which prioritize a limited number of features in line with prior biological studies. This not only increases predictive power, but also facilitates biological translation of the results as well as development of robust and simplified assays for resource-limited settings. From a bayesian perspective this could be viewed as a shift in the prior distributions over β towards estimates that are more congruent with the true underlying distributions; a more explicit connection to the bayesian setting can be seen in Methods section.

This study has several limitations, which guide our future research directions. Firstly, definition of the prior knowledge tensor by individual human experts has the potential to induce a source of bias into the analysis. While our analysis suggested that the method is robust to potential errors in the prior knowledge tensor, a more accurate and consistent definition of prior knowledge would improve this pipeline. In addition, the development of the prior knowledge is labor intensive and requires careful and objective analysis of a broad range of studies. We believe stronger results can be achieved using text-mining strategies for direct extraction of prior knowledge from the literature (*e.g.* see immuneXpresso^54^). Second, this work relies on manual analysis for identification of all cell types (Supplemental Fig. 2) and mapping them to the prior knowledge. This process is labor intensive, error-prone, and may not identify all cell populations of interest^55^. In our future studies, we will combine state-of-the-art cell population identification algorithms^56–60^ with our prior knowledge integrated to dynamically match clusters to the prior knowledge tensors for a more unbiased analysis. Third, this work only investigated incorporation of prior knowledge into the EN algorithm. However other methods can similarly be extended to incorporate expert knowledge (*e.g.*, see Krupka *et al*^*17*^. for a relevant extension of support vector machines). In our future work, we will particularly focus on incorporation of prior knowledge into deep learning methods. While these algorithms can model complex relationships that are valuable in high-throughput characterization of the immune system, the number of patients that are required for training a large neural network is often beyond the reach of typical immunological studies. We believe incorporation of prior immunological knowledge can reduce the number of patients required for implementation of deep learning approaches in clinical studies^61^. Additional research directions include: application of the iEN to domains outside of clinical immunology such as proteomics, metabolomics, and transcriptomics; and application of domain-knowledge integrated models to multi-omic studies, which would provide a systems-level perspective on human biology^62^.

## 5. Methods

### 5.1 Integration of Immunological Priors (extened)

The immunological Elastic-Net framework extends the Elastic-Net regularized regression method by integrating prior biological knowledge of cellular signal transduction into the coefficient optimization process. Consider an analysis with mass cytometry generated features *X*, composed of observations (rows) *X*^*i*^ = (*x*_*i*1_, …, *x*_*ip*_)^*T*^; for *i* = 1, 2, …, *n*, each observation consists of *p* measurements, where *p* ∈ *N* and *p* is much greater than *n*. Corresponding to each observation is a value of interest *y*_*i*_. Values of interest then constitute the response vector *Y* = (*y*_1_, …, *y*_*n*_)^*T*^. Response vectors are dataset specific (*e.g.*, a vector of gestational age during pregnancy in the LTP example). A multivariate regression model can be constructed by computing the coefficients *β* = (*β*_1_, *β*_2_, …, *β*_*p*_)^*T*^ that optimize the objective function, *L*(*β*) = ‖*Y* − *Xβ*‖^2^. The EN method expands this definition with a linear combination of two regularization terms, the *L*_1_ = ‖*β*‖_1_ and *L*_2_ = ‖*β*‖^2^ penalties, from Least Absolute Shrinkage and Selection Operator (LASSO) and Ridge regression respectively^63^. The *L*_1_ penalization reduces model complexity and increases sparsity while simultaneously selecting more descriptive features. However, it can select, at most, the number of observations when working in a high dimensional space (specifically high dimensional small observation size), and cannot select multiple, highly correlated features. *L*_2_ penalization reduces coefficient size and encourages the inclusion of highly correlated features but cannot remove features completely. Incorporating both *L*_1_ and *L*_2_ regularization terms compensates for these issues. Penalization is applied to coefficients during model fitting and is determined by a penalization factor *λ*, as well as the ratio of penalization applied to each penalty term, *α*. The optimal ratio (*α*) and degree (*λ*) of penalization can be determined through optimization of the EN objective function:

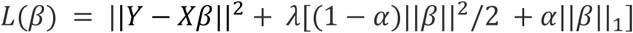

EN models are agnostic to any information not included in *X*. Whereas the iEN incorporates a third parameter which encodes prior immunological knowledge: *ϕ*, a *p* × *p* diagonal matrix of the form diag(*ϕ*)= {*φ*_1,1_, *φ*_2,2_, …, *φ*_*p,p*_} where *φ*_*i,j*_ = 0 ∀_*i*≠*j*_.The *ϕ* factor guides models to be more consistent with the current understanding of signal transduction response. Biological priors compiled *a priori* by an independent panel of immunologists is used to prioritize certain signal transduction responses via scaling features of *X*. The adapted model takes the form:

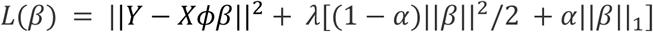

The biological priors represented as a tensor of domain specific knowledge manually constructed by a panel of experts. These biological priors are represented as a tensor of scores where features more consistent with known biology have higher values. These priors are a conservative indication of response from canonical signaling pathways that the field has a high level of confidence in observing. They are constructed as a *m* by *l* by *o* tensor, *Z* ∈ [0,1]^*m*×*l*×*o*^, where the associated mass cytometry assay consists of *m* cell types, *l* stimulations, and *o* measured responses. An element in this tensor would correspond to a particular celltype and whether it will elicit a specific signaling response in response to each *ex vivo* stimulation. To make the connection between the prior tensor *Z* and the iEN parameter *ϕ* clear, consider the function *F*(*Z*) → *diag*(*ϕ*) ∈ *R*^*P*^_>0_ that is to say, *diag*(*ϕ*) is a vector of dimension *m* · *l* · *o* = *p* that exists within the *p*-dimensional positive real numbers. That is, *Z* is transformed to a *p*-dimensional vector, where *diag*(*ϕ*) = {*φ*_1,1_, *φ*_2,2_, …, *φ*_*p,p*_}such that 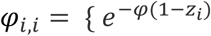where *z*_*i*_ is the score of the *i*^*th*^ immune feature, and *φ* is the amount of prioritization applied}. This formulation of *φ*_*i,i*_ ∈ *diag*(*ϕ*) as 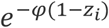 affects features with lower prior value more so than features with larger values. This definition allows for increased model stability than a formulation with *φ*_*i,i*_ ∈ *diag*(*ϕ*) as 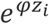 for large *φ*′*s* (Supplemental Fig. 1a/b).

### 5.2 Bayesian Interpretation

The Elastic-Net has a Bayesian representation^64^ with priors over the estimates of *β*. This can help define the role of immunological priors in increasing predictive power. The unnormalized version of this prior is reported as follows:

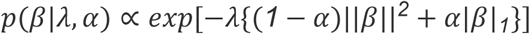

In the following, we show that the immunological Elastic-Net has a similar interpretation in which the prior distributions over *β* are altered according to the prioritization of biological knowledge, i.e. the value of *φ* and the shape of *Z*. That is to say, our definition of the iEN can be represented as an alteration of the prior distribution over *β* given *ϕ*. The objective function of the iEN is as follows, 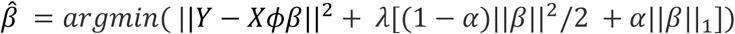. Now let 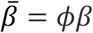, substitution for *β* results in the following optimization problem:

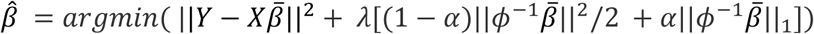

From this formulation, the adjusted Bayesian prior for the iEN can be directly derived as follows:

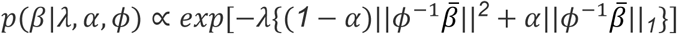

To further illustrate the connection between iEN and the regular EN and their Bayesian interpretations, we show that EN is a special case of iEN. For this, let us define the following two sets:

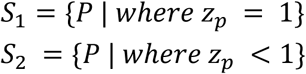

These sets indicate which estimates of 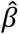 are affected by *φ* and which remain unaffected as previously defined. We can then subset the *Z* vector accordingly with 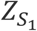 being all biological priors of value one and 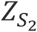 being all biological priors of value less than one. Therefore, we can separate the *L*_1_and *L*_2_norms according to these sets reformulating the optimization problem as follows:

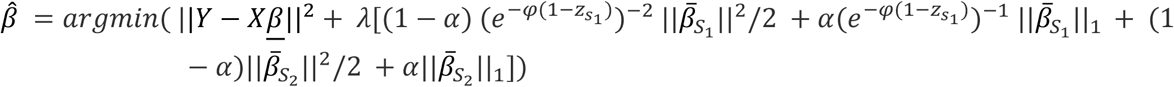

Here 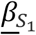 and 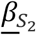 represent the betas which are affected by the *φ* value. This allows for us to replace *ϕ* with *e*^−*φ*(1−*z*)^ respective of *S*_1_and *S*_2_, which demonstrates how prioritization affects the estimation of *β*. Similar separation in the prior distribution also illustrates how priors over *β* are affected in the same manner.

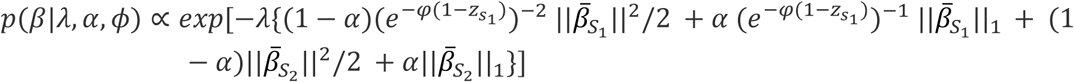

Since *φ* alters the prior distribution over *β*, this can be used to improve estimates of the true *β*in the iEN’s Bayesian setting, as it does in the classic formulation. This also allows for the EN estimates to be included as a special case of the iEN when *φ* = 0.

### 5.3 Parameter Optimization

iEN parameters were optimized over a grid of possible parameter values for each parameter (*φ*, *α, λ*). The *φ* search grid was generated from a logarithmic sequence with values between zero and 100, which allows for the EN as a special case (*φ*= 0). Similarly, *α* was uniformly generated from values between zero and one. Generation of *λ* was done so that a large range of model sizes were tested during the 10-fold CV. Specifically, *λ* values were generated during the inner CV loop to avoid any possible information leak. Metrics used to justify parameter selection were residual sum of squares for continuous response and area under the receiver operating characteristic curve, as appropriate for each example presented.

### 5.4 Simulated Data

All simulated data were generated using the *‘simglm*’ R package^65^. 450 features were generated with a standard deviation of 15. However, 200 of these features had a mean value sampled from 𝒩(𝒩(0, 10^2^), 15^2^) to simulate features moderately associated with prior knowledge and 50 were sampled from 𝒩(𝒩(0, 10^2^), 15^2^), representing features highly associated with prior knowledge. The response variable is then generated as a linear combination of these 250 features. The additional 450 features represented features not associated with prior knowledge and were generated randomly using a normal distribution.

### 5.5 LTP Cohort

Twenty-one pregnant women were included in the LTP study, all of whom received routine antepartum care at Lucile Packard Children’s Hospital. Three patients were excluded from the study due to premature delivery (<37 weeks of gestation). Analysis was performed on the remaining eighteen women who delivered at term (≥37 weeks of gestation). These eighteen participants were age 31.9 (years) ± 3.4 (standard deviation) old. An independent cohort of 10 pregnant women who delivered at term were later enrolled as a validation cohort.

### 5.6 ChP Cohort

A total of thirty patients were enrolled in the study of ChP, 15 healthy controls and 15 patients with ChP receiving treatment at Bell Dental Center (San Leandro, CA) and Stanford University School of Medicine (Stanford, CA). Two participants were excluded from the analysis, one patient due to autoimmune disease and one control due to onset of hand infection during the study. The final cohort consisted of 14 patients (age 42.2 ± 10.5) and 14 controls (age 36.5 ± 8.07) samples, each of which were split by gender: eight female, six male.

### 5.7 Whole Blood Sampling

Whole blood samples were collected in 10mL heparin-containing tubes and processed within one hour of collection. Samples for the LTP cohort were stimulated with either 1ug/mL of Lipopolysaccharide (LPS), 100ng/mL of Interferon-*α* (IFN*α*), or a cocktail of 100ng/mL of Interleukins (IL-2, IL-6), or they were left unstimulated to measure endogenous cellular activity. Samples for the ChP cohort were stimulated with LPS, IFN*α*, TNF*α*, or a cocktail of IL-2, IL-4, IL-6 and GM-CSF or left unstimulated. Samples were fixed using a stabilization buffer (SmartTube Inc.) according to manufacturer instructions and stored at -80°C until further processing.

### 5.8 Mass Cytometry Analysis

Post-thaw, fixed samples were added to an erythrocyte lysis buffer (SmartTube Inc) and underwent two rounds of erythrocyte lysis. Cells were then barcoded as previously described^66^. In summary, twenty-well barcode plates were prepared with a combination of 2 Pd isotopes out of a pool of six (^102^Pd, ^104^Pd, ^105^Pd, ^106^Pd, ^108^Pd, ^110^Pd) and added to the cells in 0.02% saponin/PBS. Samples were pooled and stained with metal-conjugated antibodies collectively to minimize experimental variation. The panel for the different cohorts are listed in Supplementary tables 3, and 4. Intracellular staining was performed in methanol-permeabilized cells. Cells were incubated overnight at 4°C with an iridium-containing intercalator (Fluidigm). Prior to mass cytometry analysis, cells were filtered through a 35μm membrane and resuspended in a solution of normalization beads (Fluidigm).

Barcoded and stained cells were analysed on a Helios Mass Cytometer (Fluidigm) at an event rate of 500 to 1000 cells per second. The data was normalized using Normalizer v0.1 MATLAB Compiler Runtime^67^ and debarcoded with a single-cell MATLAB debarcoding tool^66^. Gating was performed using Cytobank (cytobank.org). Gating strategies for the different cohorts are shown in Supplementary Fig. 4.

