## supplemental materials for "Integration of Mechanistic Immunological Knowledge into a Machine Learning Pipeline Increases Predictive Power"

### Supplemental Information

#### Supplemental Figures:

**a.** $e^{\varphi z_i}$ Shrinkage

**b.** $e^{-\varphi(1-z_i)}$ Shrinkage

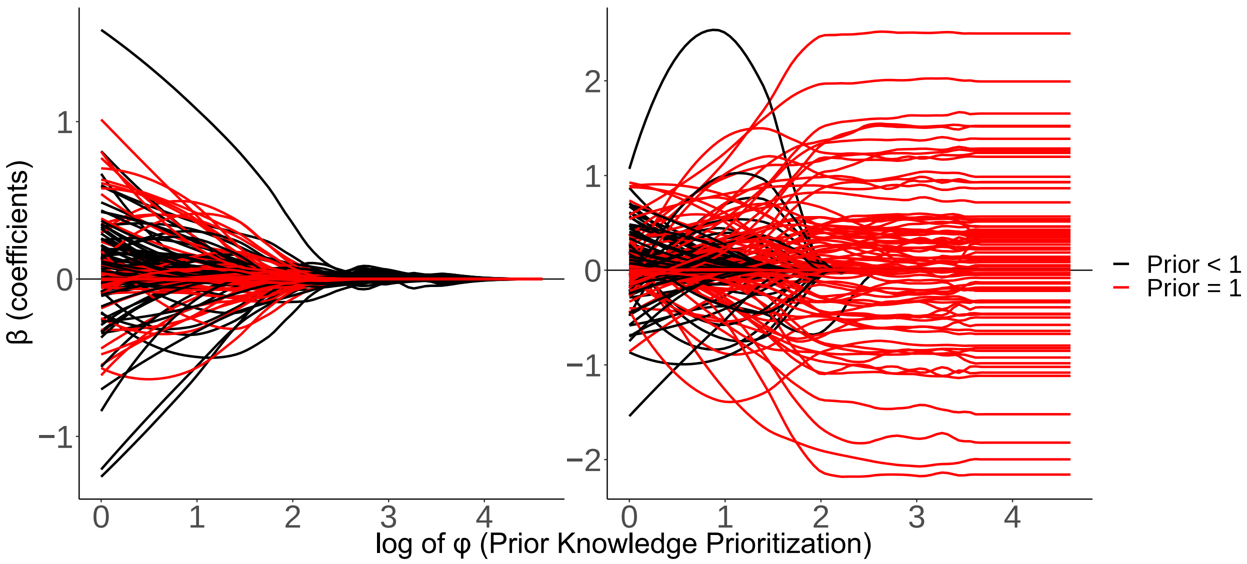


**c.** Model Sparcification

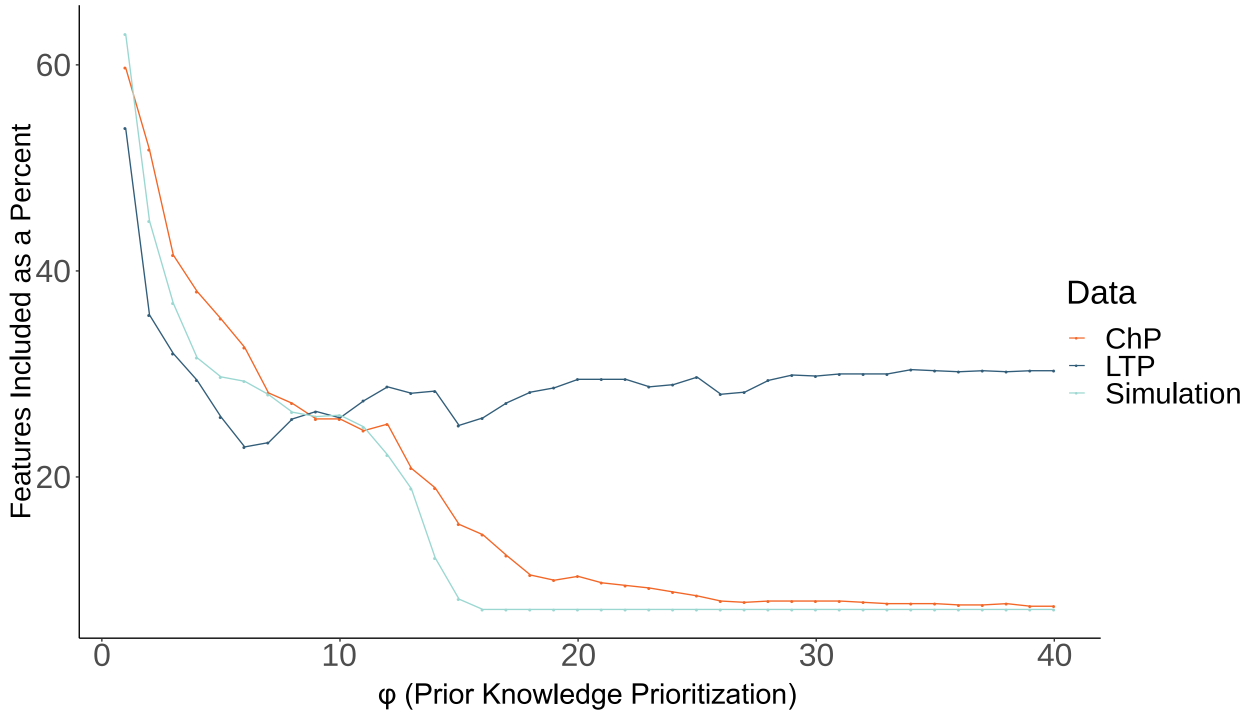

**[Supplemental Figure 1 - Effect of feature prioritization on coefficients and model size: (a)** prior knowledge prioritization with $\varphi_{i,i} \in diag(\phi)$ as $e^{\varphi z_i}$ results in instability. This holds true for features with both high and low prior knowledge values. This instability is rooted in values truncating to infinity as $\phi$ increases, effectively removing all features in the order of highest prior value to lowest prior value. **(b)** Prioritization with $\varphi_{i,i} \in diag(\phi)$ as $e^{-\varphi(1-z_i)}$ results in stable progression of feature shrinkage as prioritization increases. This allows for the edge case where the model is constructed solely from features with prior value = 1. **(c)** Variation of $\varphi$ to a model with fixed $\alpha$ and $\lambda$ ($\alpha$ = 0.5, with $\lambda$ selected such that each initial dataset had comparable model size) affects the number of features included, presented here as percent of features for each dataset. An increase of $\varphi$ in all datasets follow similar trends, note that the ChP and Simulated studies truncate at approximately 7% of features included. This suggests that the simulated study reacts similarly to real mass cytometry data for evaluation of the model optimization procedure.]

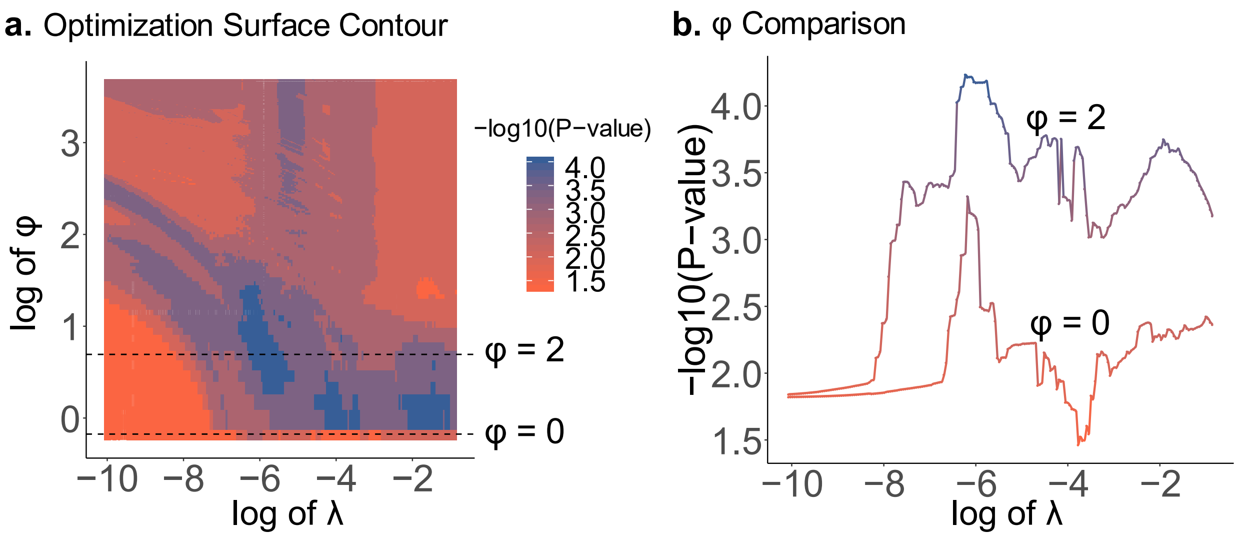


[**Supplemental Figure 2 - Optimization landscape**: **(a)** Optimization of iEN models for the LTP study across a grid of $\varphi$ and $\lambda$ values (with $\alpha$ = 0.5) demonstrates the improvement in performance after incorporation of prior knowledge ($\varphi$ = 0 is equivalent to the EN). The color gradient of this surface represents the -log10(P-value) of models trained on the original LTP data which predict the Validation dataset, for combinations of $\lambda \in$ [4.25e-5,0.425] and $\varphi \in$ [0,40]. This indicates that models fit with prioritization of expert knowledge ($\varphi$ = 2) outperform the traditional EN ($\varphi$ = 0). **(b)** A direct comparison of models, EN ($\varphi$ = 0) vs. iEN ($\varphi$ = 2) as indicated by the dashed lines in panel a, visualizing the performance difference over the $\lambda$ sequence. This clearly demonstrates a robust improvement in the results when prior knowledge is integrated into the model.]

**a.** Extended Simulation Study

**b.** Runtime Analysis

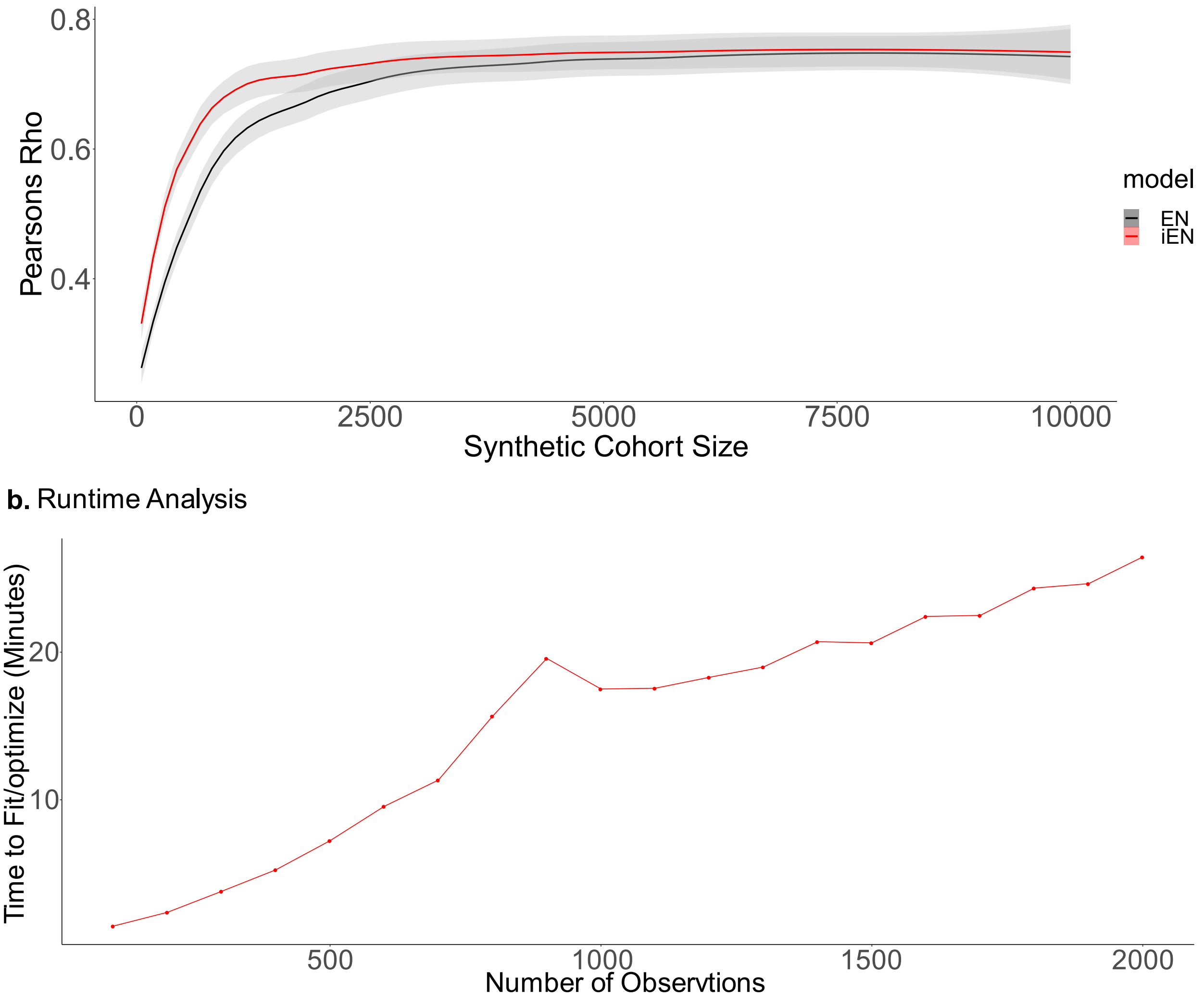


[**Supplemental Figure 3 - Extended results from the simulation study:: (a)** Simulated data was used to compare iEN and EN across a broad range of cohort sizes. Twenty simulated cohorts ranging from 50 to 10000 subjects (selected from a logarithmic sequence) were used to generate these datasets. Each cohort of simulated "patients" were fit repeatedly 10 times using a randomized 10-fold cross-validation stratification. iEN outperformed EN in cohorts as large as 2000 patients. **(b)** The simulated data was also used to determine the time requirement of the iEN algorithm with varying population sizes (100 to 2000). This demonstrated a linear relationship between the number of observations within the data and the time to completion. Each model was optimized using the same parameter search space and 10-fold CV strategy used in the clinical studies. Here computation time for only one iteration is reported because the iEN software used is parallelized over each combination of $\varphi$ and $\alpha$ values for high-performance computing.**]**

### Gating Strategy

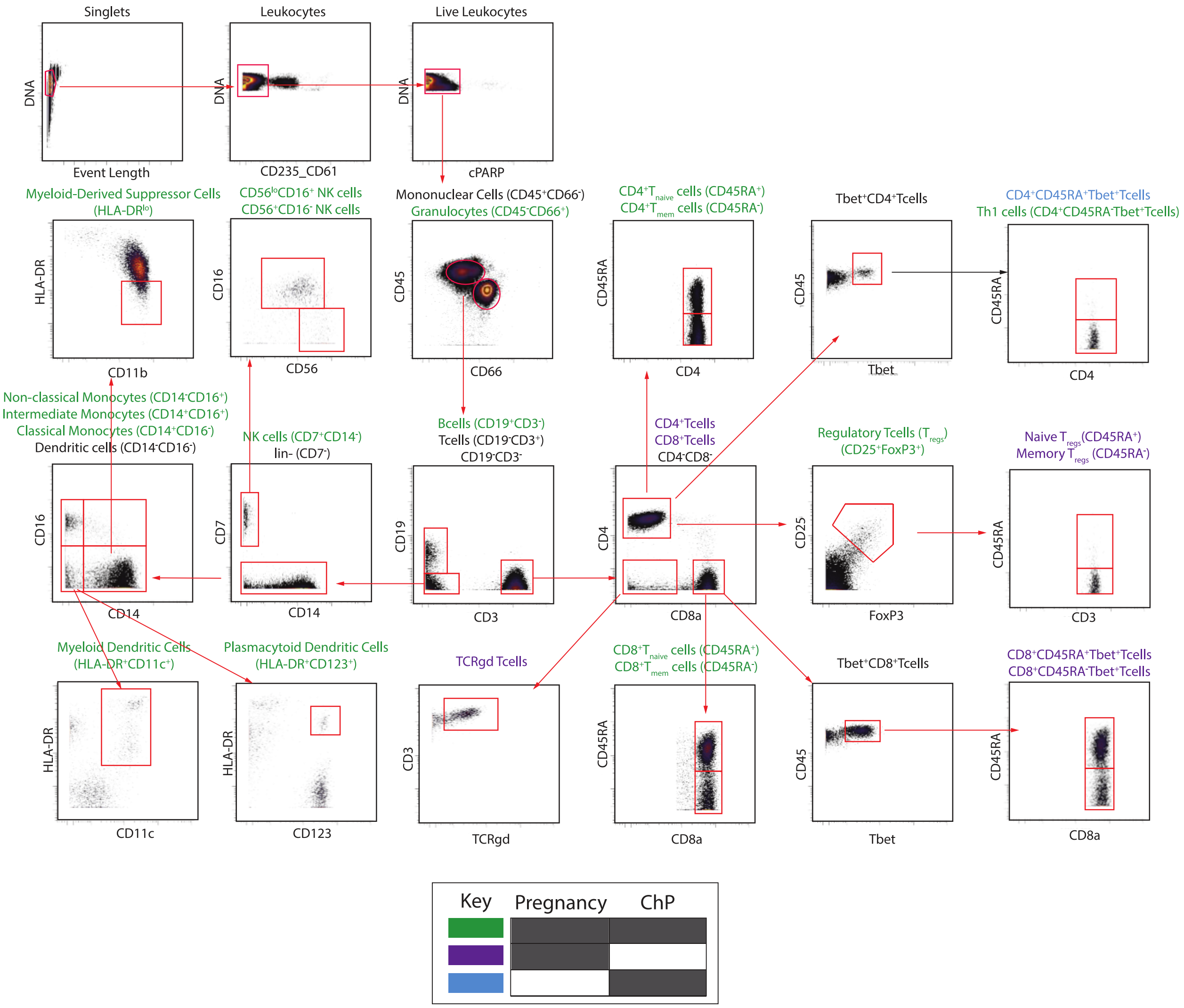


[**Supplemental Figure 4 - Gating strategy for feature extraction from the measured single cells (combined for both clinical datasets):** Two-dimensional scatter plots shown for a representative patient sample. Gating was performed using Cytobank (www.cytobank.org) for the ChP and LTP cohorts. Populations noted in green were included in both analyses, purple populations were included in LTP analysis and blue populations were supplemented in only the ChP analysis.]

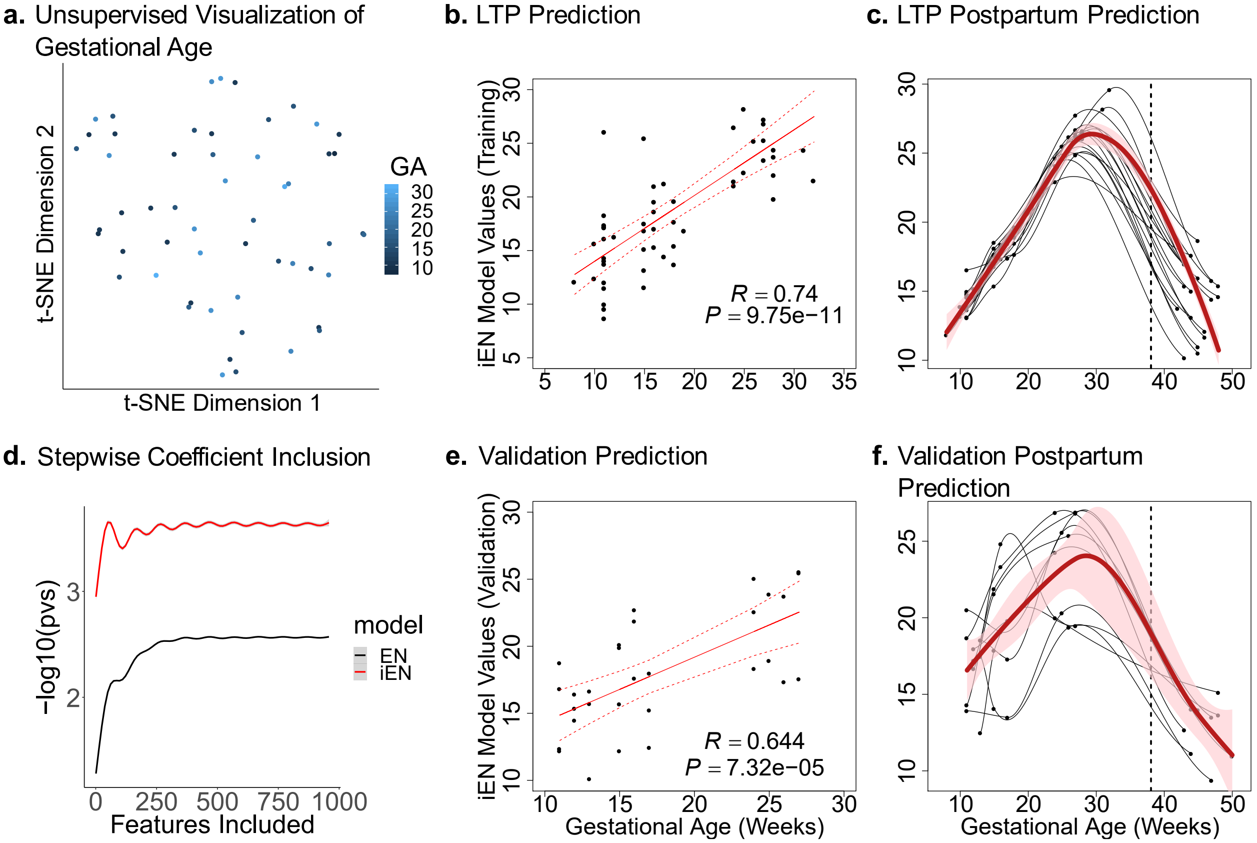


[**Supplemental Figure 5 - Analysis of longitudinal term pregnancy**: **(a)** t-SNE visualization of each antepartum sample colored by gestational age shows no major connection between the measured features and gestational age. **(b)** Mean prediction from 10-fold cross-validated iEN models on training data shows the strong correlation of this knowledge integrated model's estimations with GA. **(c)** Projecting mean coefficient models onto training data which contains postpartum samples demonstrates a return to baseline after delivery. **(d)** Smoothing splines of Pearson correlation p-value generated from each individual iEN and EN model, applied to the independent validation cohort with descending stepwise inclusion of features by coefficient size. **(e)** Predicted vs. actual gestational age values and correlation test results. **(f)** Similar to the training cohort, models applied to the validation cohort demonstrate a postpartum return to baseline.]

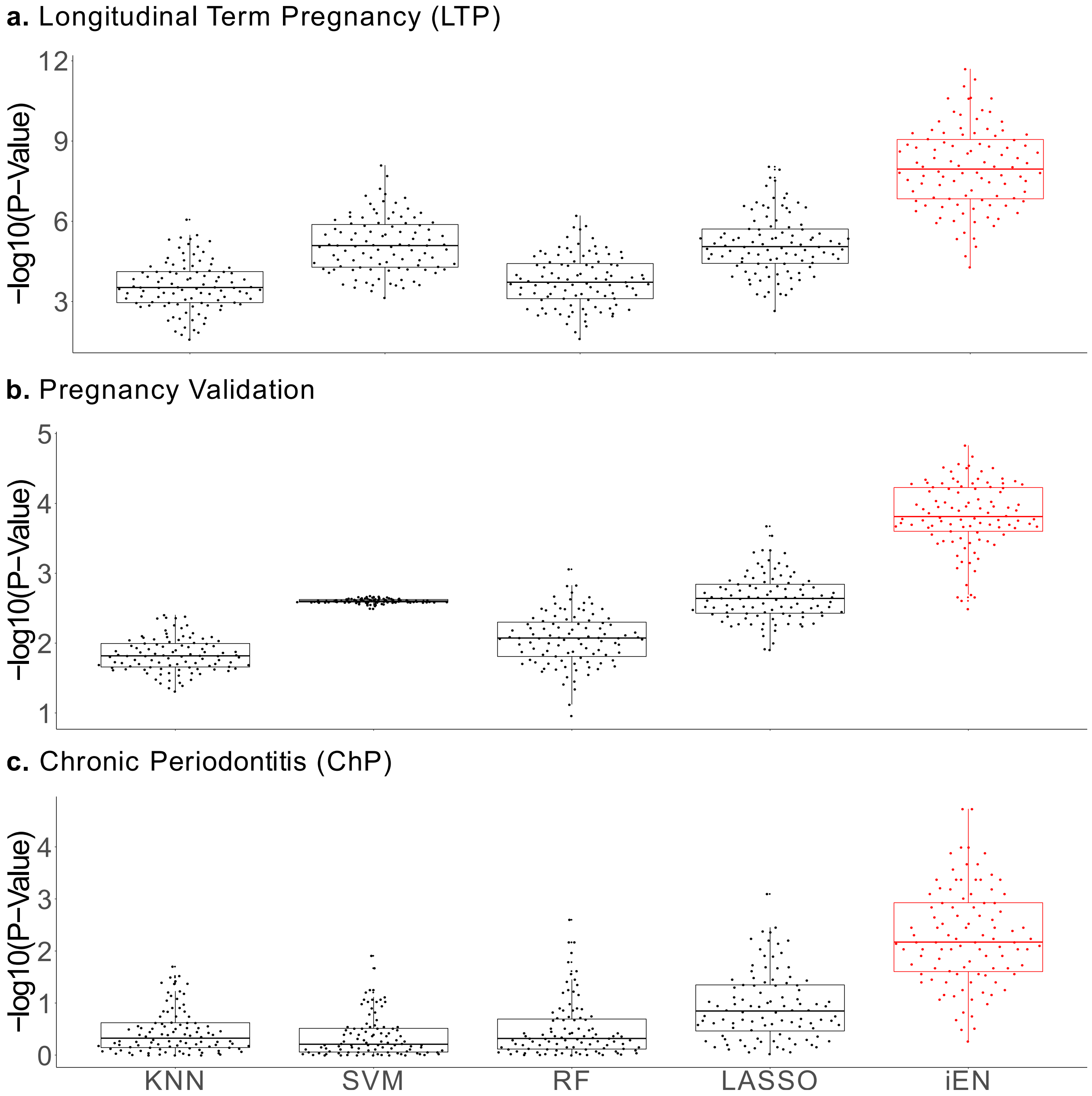


[**Supplemental Figure 6 - Comparison against standard machine learning algorithms:** Expanded comparison of the IEN method against standard machine learning algorithms for **(a)** LTP**, (b)** LTP validation, and **(c)** ChP analysis. All algorithms were trained, tested, and optimized using a similar nested 10-fold CV approach.]

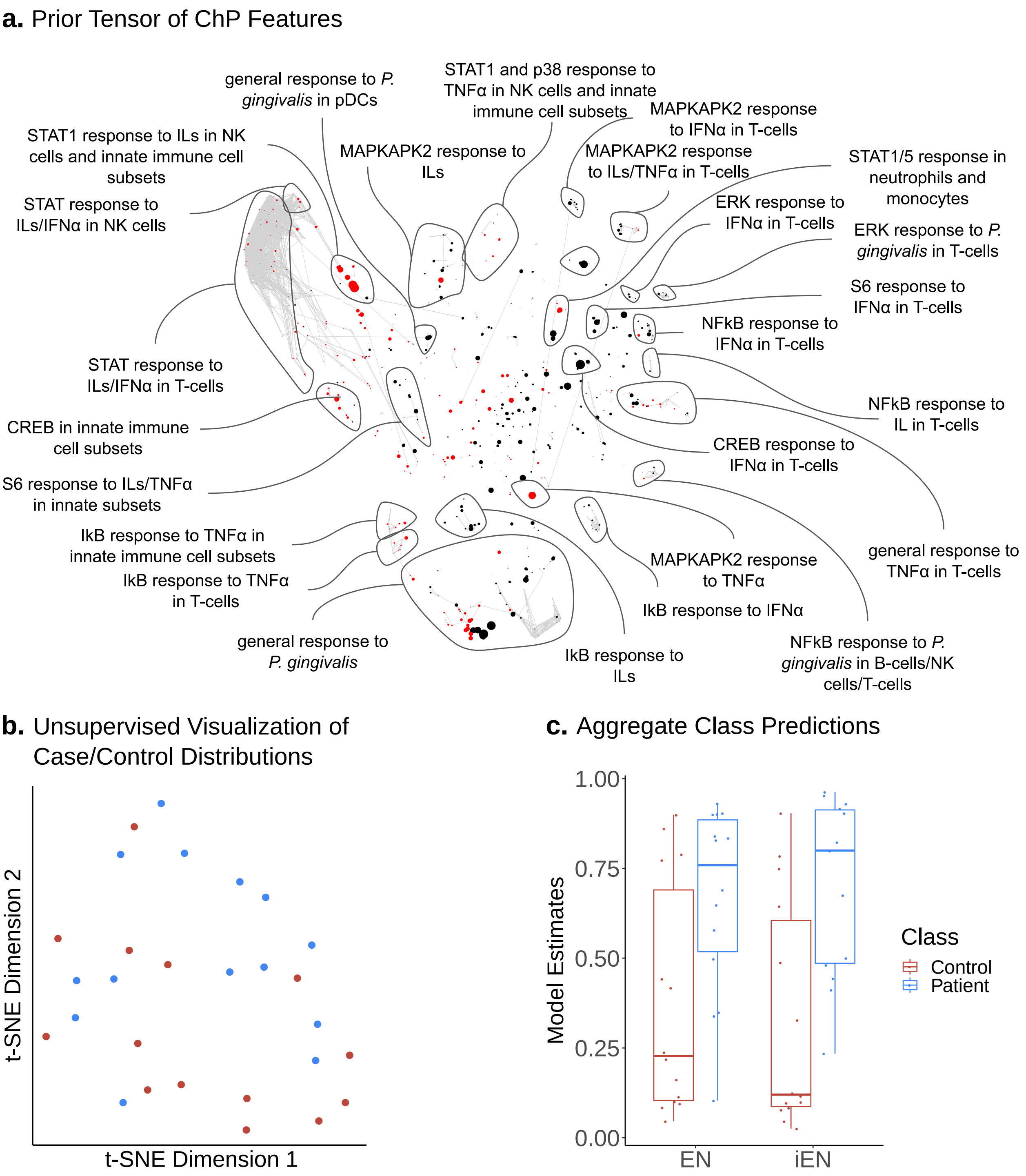


[**Supplemental Figure 7 - Analysis of chronic periodontitis**: **(a)** A correlation network of intracellular signaling responses, measured in peripheral immune cells with nodes representing canonical signaling pathways (i.e. indicated by biological priors with a score > 0.5) displayed in red, otherwise black. Edges represent significant (P-value < 0.05) pairwise correlation after Bonferroni correction. Node size represents the significance of correlation to the response variable. **(b)** t-SNE visualization of patients shows no clear separation between control and patient populations across all immune features, making this a suitable setting for supervised classification analysis. **(c)** Boxplot of mean out-of-sample prediction from 10-fold CV. Access to prior knowledge improved iEN's results compared to that of EN (with no access to prior knowledge).]

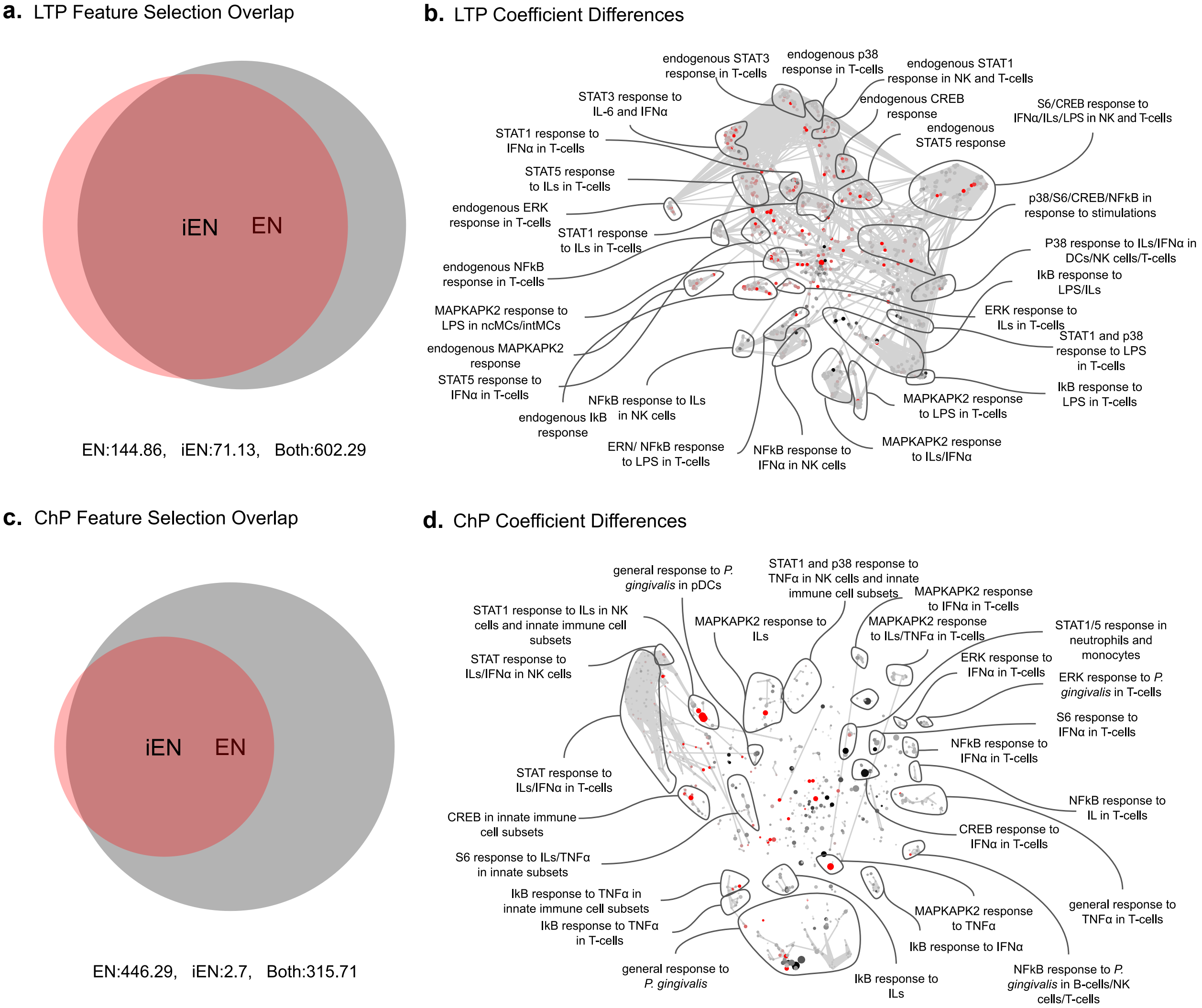


[**Supplemental Figure 8 - iEN and EN feature selection and coefficient comparison: (a)** Comparison of features selected between each cross-validation iteration for LTP analysis shows significant overlap between the two models with slightly larger EN models. **(b)** Further comparison of iEN and EN LTP model coefficients averaged across all cross-validation iterations displays areas of the feature space more represented in their respective model. Immune features which are red are more represented in iEN models, while black nodes are more represented in EN models. **(c)** Comparison of features selected during ChP analysis shows similar overlap of features selected. However, EN models consistently used a larger number of features for the ChP than the LTP analysis. **(d)** ChP coefficients between iEN and EN models also displays a difference in feature representation.]

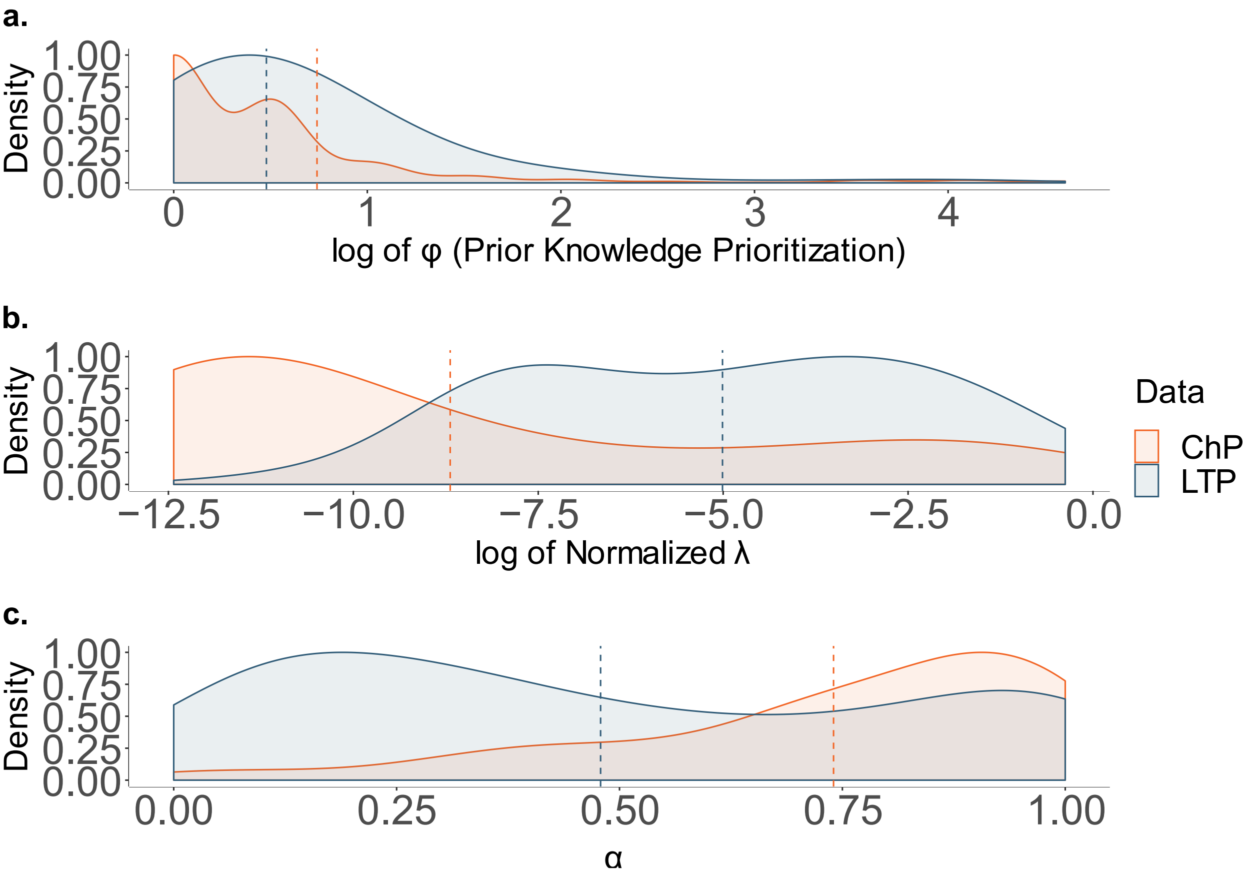


[**Supplemental Figure 9 - Parameter selection during cross-validation:** Scaled distribution of optimized parameters, **(a)** $\varphi$, **(b)** $\lambda$, and **(c)** $\alpha$, colored by dataset. The mean of each distribution is overlaid as a dashed line. The consistency of parameter optimization across datasets and parameters show the stability of the model during the optimization process. Both $\varphi$ and $\lambda$ were generated on a log scale and are visualized accordingly.]

#### Supplemental Tables:

**Table 1. Prior knowledge matrix for estimation of gestational age.**

| **Endogenous** | pCREB | pERK1/2 | IkB | pMK2 | pNFkB | pP38 | pS6 | pSTAT1 | pSTAT3 | pSTAT5 |
|---|---|---|---|---|---|---|---|---|---|---|
| Bcells | 1 | 1 | 1 | 1 | 1 | 1 | 1 | 1 | 1 | 1 |
| $CD16^{+}CD56^{-}$NKcells | 1 | 1 | 1 | 1 | 1 | 1 | 1 | 1 | 1 | 1 |
| $CD4^{+}Tcells_{mem}$ | 1 | 1 | 1 | 1 | 1 | 1 | 1 | 1 | 1 | 1 |
| $CD4^{+}Tcells_{naive}$ | 1 | 1 | 1 | 1 | 1 | 1 | 1 | 1 | 1 | 1 |
| $CD4^{+}$Tcells | 1 | 1 | 1 | 1 | 1 | 1 | 1 | 1 | 1 | 1 |
| $CD45RA^{+}$Tregs | 1 | 1 | 1 | 1 | 1 | 1 | 1 | 1 | 1 | 1 |
| $CD45RA^{-}$Tregs | 1 | 1 | 1 | 1 | 1 | 1 | 1 | 1 | 1 | 1 |
| $CD56^{+}CD16^{-}$NKcells | 1 | 1 | 1 | 1 | 1 | 1 | 1 | 1 | 1 | 1 |
| $CD7^{+}$NKcells | 1 | 1 | 1 | 1 | 1 | 1 | 1 | 1 | 1 | 1 |
| $CD8^{+}Tcells_{mem}$ | 1 | 1 | 1 | 1 | 1 | 1 | 1 | 1 | 1 | 1 |
| $CD8^{+}Tcells_{naive}$ | 1 | 1 | 1 | 1 | 1 | 1 | 1 | 1 | 1 | 1 |
| $CD8^{+}$Tcells | 1 | 1 | 1 | 1 | 1 | 1 | 1 | 1 | 1 | 1 |
| cMCs | 1 | 1 | 1 | 1 | 1 | 1 | 1 | 1 | 1 | 1 |
| Granulocytes | 1 | 1 | 1 | 1 | 1 | 1 | 1 | 1 | 1 | 1 |
| intMCs | 1 | 1 | 1 | 1 | 1 | 1 | 1 | 1 | 1 | 1 |
| M-MDSC | 1 | 1 | 1 | 1 | 1 | 1 | 1 | 1 | 1 | 1 |
| mDCs | 1 | 1 | 1 | 1 | 1 | 1 | 1 | 1 | 1 | 1 |
| ncMCs | 1 | 1 | 1 | 1 | 1 | 1 | 1 | 1 | 1 | 1 |
| pDCs | 1 | 1 | 1 | 1 | 1 | 1 | 1 | 1 | 1 | 1 |
| $Tbet^{+}CD4^{+}Tcells_{mem}$ | 1 | 1 | 1 | 1 | 1 | 1 | 1 | 1 | 1 | 1 |
| $Tbet^{+}CD8^{+}Tcells_{mem}$ | 1 | 1 | 1 | 1 | 1 | 1 | 1 | 1 | 1 | 1 |
| $Tbet^{+}CD8^{+}Tcells_{naive}$ | 1 | 1 | 1 | 1 | 1 | 1 | 1 | 1 | 1 | 1 |
| $TCR\gamma\delta^{+}$Tcells | 1 | 1 | 1 | 1 | 1 | 1 | 1 | 1 | 1 | 1 |
| Tregs | 1 | 1 | 1 | 1 | 1 | 1 | 1 | 1 | 1 | 1 |

| IFNα | pCREB | pERK1/2 | IkB | pMK2 | pNFkB | pP38 | pS6 | pSTAT1 | pSTAT3 | pSTAT5 |
|---|---|---|---|---|---|---|---|---|---|---|
| Bcells | 0.4 | 0.4 | 0.4 | 0.4 | 0.4 | 0.4 | 0.2 | 1 | 0.8 | 0.7 |
| $CD16^{+}CD56^{-}$NKcells | 0.4 | 0.4 | 0.4 | 0.4 | 0.4 | 0.4 | 0.2 | 1 | 0.8 | 0.7 |
| $CD4^{+}Tcells_{mem}$ | 0.4 | 0.4 | 0.4 | 0.4 | 0.4 | 0.4 | 0.2 | 1 | 0.8 | 0.7 |
| $CD4^{+}Tcells_{naive}$ | 0.4 | 0.4 | 0.4 | 0.4 | 0.4 | 0.4 | 0.2 | 1 | 0.8 | 0.7 |
| $CD4^{+}$Tcells | 0.4 | 0.4 | 0.4 | 0.4 | 0.4 | 0.4 | 0.2 | 1 | 0.8 | 0.7 |
| $CD45RA^{+}$Tregs | 0.4 | 0.4 | 0.4 | 0.4 | 0.4 | 0.4 | 0.2 | 1 | 0.8 | 0.7 |
| $CD45RA^{-}$Tregs | 0.4 | 0.4 | 0.4 | 0.4 | 0.4 | 0.4 | 0.2 | 1 | 0.8 | 0.7 |
| $CD56^{+}CD16^{-}$NKcells | 0.4 | 0.4 | 0.4 | 0.4 | 0.4 | 0.4 | 0.2 | 1 | 0.8 | 0.7 |
| $CD7^{+}$NKcells | 0.4 | 0.4 | 0.4 | 0.4 | 0.4 | 0.4 | 0.2 | 1 | 0.8 | 0.7 |
| $CD8^{+}Tcells_{mem}$ | 0.4 | 0.4 | 0.4 | 0.4 | 0.4 | 0.4 | 0.2 | 1 | 0.8 | 0.7 |
| $CD8^{+}Tcells_{naive}$ | 0.4 | 0.4 | 0.4 | 0.4 | 0.4 | 0.4 | 0.2 | 1 | 0.8 | 0.7 |
| $CD8^{+}$Tcells | 0.4 | 0.4 | 0.4 | 0.4 | 0.4 | 0.4 | 0.2 | 1 | 0.8 | 0.7 |

| | | | | | | | | | | |
|---|---|---|---|---|---|---|---|---|---|---|
| cMCs | 0.4 | 0.4 | 0.4 | 0.4 | 0.4 | 0.4 | 0.2 | 1 | 0.8 | 0.7 |
| Granulocytes | 0.4 | 0.4 | 0.4 | 0.4 | 0.4 | 0.4 | 0.2 | 1 | 0.8 | 0.7 |
| intMCs | 0.4 | 0.4 | 0.4 | 0.4 | 0.4 | 0.4 | 0.2 | 1 | 0.8 | 0.7 |
| M-MDSC | 0.4 | 0.4 | 0.4 | 0.4 | 0.4 | 0.4 | 0.2 | 1 | 0.8 | 0.7 |
| mDCs | 0.4 | 0.4 | 0.4 | 0.4 | 0.4 | 0.4 | 0.2 | 1 | 0.8 | 0.7 |
| ncMCs | 0.4 | 0.4 | 0.4 | 0.4 | 0.4 | 0.4 | 0.2 | 1 | 0.8 | 0.7 |
| pDCs | 0.4 | 0.4 | 0.4 | 0.4 | 0.4 | 0.4 | 0.2 | 1 | 0.8 | 0.7 |
| Tbet$^+$CD4$^+$Tcells$_{mem}$ | 0.4 | 0.4 | 0.4 | 0.4 | 0.4 | 0.4 | 0.2 | 1 | 0.8 | 0.7 |
| Tbet$^+$CD8$^+$Tcells$_{mem}$ | 0.4 | 0.4 | 0.4 | 0.4 | 0.4 | 0.4 | 0.2 | 1 | 0.8 | 0.7 |
| Tbet$^+$CD8$^+$Tcells$_{naive}$ | 0.4 | 0.4 | 0.4 | 0.4 | 0.4 | 0.4 | 0.2 | 1 | 0.8 | 0.7 |
| TCR$\gamma\delta^+$Tcells | 0.4 | 0.4 | 0.4 | 0.4 | 0.4 | 0.4 | 0.2 | 1 | 0.8 | 0.7 |
| Tregs | 0.4 | 0.4 | 0.4 | 0.4 | 0.4 | 0.4 | 0.2 | 1 | 0.8 | 0.7 |

| IL (IL-2/IL-6) | pCREB | pERK1/2 | IkB | pMK2 | pNFkB | pP38 | pS6 | pSTAT1 | pSTAT3 | pSTAT5 |
|---|---|---|---|---|---|---|---|---|---|---|
| Bcells | 0.3 | 0.6 | 0.3 | 0.2 | 0.3 | 0.2 | 0.2 | 0.7 | 0.8 | 0.8 |
| CD16$^+$CD56$^-$NKcells | 0.3 | 0.6 | 0.3 | 0.2 | 0.3 | 0.2 | 0.2 | 0.7 | 0.8 | 0.9 |
| CD4$^+$Tcells$_{mem}$ | 0.3 | 0.6 | 0.3 | 0.2 | 0.3 | 0.2 | 0.2 | 0.7 | 0.8 | 0.9 |
| CD4$^+$Tcells$_{naive}$ | 0.3 | 0.6 | 0.3 | 0.2 | 0.3 | 0.2 | 0.2 | 0.7 | 0.8 | 0.9 |
| CD4$^+$Tcells | 0.3 | 0.6 | 0.3 | 0.2 | 0.3 | 0.2 | 0.2 | 0.7 | 0.8 | 0.9 |
| CD45RA$^+$Tregs | 0.3 | 0.6 | 0.3 | 0.2 | 0.3 | 0.2 | 0.2 | 0.7 | 0.8 | 0.9 |
| CD45RA$^-$Tregs | 0.3 | 0.6 | 0.3 | 0.2 | 0.3 | 0.2 | 0.2 | 0.7 | 0.8 | 0.9 |
| CD56$^+$CD16$^-$NKcells | 0.3 | 0.6 | 0.3 | 0.2 | 0.3 | 0.2 | 0.2 | 0.7 | 0.8 | 0.9 |
| CD7$^+$NKcells | 0.3 | 0.6 | 0.3 | 0.2 | 0.3 | 0.2 | 0.2 | 0.7 | 0.8 | 0.9 |
| CD8$^+$Tcells$_{mem}$ | 0.3 | 0.6 | 0.3 | 0.2 | 0.3 | 0.2 | 0.2 | 0.7 | 0.8 | 0.9 |
| CD8$^+$Tcells$_{naive}$ | 0.3 | 0.6 | 0.3 | 0.2 | 0.3 | 0.2 | 0.2 | 0.7 | 0.8 | 0.9 |
| CD8$^+$Tcells | 0.3 | 0.6 | 0.3 | 0.2 | 0.3 | 0.2 | 0.2 | 0.7 | 0.8 | 0.9 |
| cMCs | 0.3 | 0.6 | 0.3 | 0.2 | 0.3 | 0.2 | 0.2 | 0.6 | 0.8 | 0.7 |
| Granulocytes | 0.2 | 0.3 | 0.2 | 0.2 | 0.2 | 0.2 | 0.2 | 0.2 | 0.6 | 0.6 |
| intMCs | 0.3 | 0.6 | 0.3 | 0.2 | 0.3 | 0.2 | 0.2 | 0.6 | 0.8 | 0.7 |
| M-MDSC | 0.3 | 0.6 | 0.3 | 0.2 | 0.3 | 0.2 | 0.2 | 0.6 | 0.8 | 0.7 |
| mDCs | 0.3 | 0.6 | 0.3 | 0.2 | 0.3 | 0.2 | 0.2 | 0.6 | 0.8 | 0.7 |
| ncMCs | 0.3 | 0.6 | 0.3 | 0.2 | 0.3 | 0.2 | 0.2 | 0.6 | 0.7 | 0.7 |
| pDCs | 0.3 | 0.6 | 0.3 | 0.2 | 0.3 | 0.2 | 0.2 | 0.6 | 0.8 | 0.7 |
| Tbet$^+$CD4$^+$Tcells$_{mem}$ | 0.3 | 0.6 | 0.3 | 0.2 | 0.3 | 0.2 | 0.2 | 0.7 | 0.8 | 0.9 |
| Tbet$^+$CD8$^+$Tcells$_{mem}$ | 0.3 | 0.6 | 0.3 | 0.2 | 0.3 | 0.2 | 0.2 | 0.7 | 0.8 | 0.9 |
| Tbet$^+$CD8$^+$Tcells$_{naive}$ | 0.3 | 0.6 | 0.3 | 0.2 | 0.3 | 0.2 | 0.2 | 0.7 | 0.8 | 0.9 |
| TCR$\gamma\delta^+$Tcells | 0.3 | 0.6 | 0.3 | 0.2 | 0.3 | 0.2 | 0.2 | 0.7 | 0.8 | 0.9 |
| Tregs | 0.3 | 0.6 | 0.3 | 0.2 | 0.3 | 0.2 | 0.2 | 0.7 | 0.8 | 0.9 |

| LPS | pCREB | pERK1/2 | IkB | pMK2 | pNFkB | pP38 | pS6 | pSTAT1 | pSTAT3 | pSTAT5 |
|---|---|---|---|---|---|---|---|---|---|---|
| Bcells | 0.2 | 0.2 | 0.2 | 0.2 | 0.2 | 0.2 | 0.2 | 0 | 0 | 0 |
| $CD16^{+}CD56^{-}$NKcells | 0.7 | 0.7 | 0.7 | 0.7 | 0.7 | 0.7 | 0.7 | 0 | 0 | 0 |
| $CD4^{+}Tcells_{mem}$ | 0 | 0 | 0 | 0 | 0 | 0 | 0 | 0 | 0 | 0 |
| $CD4^{+}Tcells_{naive}$ | 0 | 0 | 0 | 0 | 0 | 0 | 0 | 0 | 0 | 0 |
| $CD4^{+}$Tcells | 0 | 0 | 0 | 0 | 0 | 0 | 0 | 0 | 0 | 0 |
| $CD45RA^{+}$Tregs | 0.2 | 0.2 | 0.2 | 0.2 | 0.2 | 0.2 | 0.2 | 0 | 0 | 0 |
| $CD45RA^{-}$Tregs | 0.2 | 0.2 | 0.2 | 0.2 | 0.2 | 0.2 | 0.2 | 0 | 0 | 0 |
| $CD56^{+}CD16^{-}$NKcells | 0.7 | 0.7 | 0.7 | 0.7 | 0.7 | 0.7 | 0.7 | 0 | 0 | 0 |
| $CD7^{+}$NKcells | 0.7 | 0.7 | 0.7 | 0.7 | 0.7 | 0.7 | 0.7 | 0 | 0 | 0 |
| $CD8^{+}Tcells_{mem}$ | 0 | 0 | 0 | 0 | 0 | 0 | 0 | 0 | 0 | 0 |
| $CD8^{+}Tcells_{naive}$ | 0 | 0 | 0 | 0 | 0 | 0 | 0 | 0 | 0 | 0 |
| $CD8^{+}$Tcells | 0 | 0 | 0 | 0 | 0 | 0 | 0 | 0 | 0 | 0 |
| cMCs | 1 | 1 | 1 | 1 | 1 | 1 | 1 | 0.2 | 0.2 | 0.2 |
| Granulocytes | 0.7 | 0.7 | 0.7 | 0.7 | 0.7 | 0.7 | 0.7 | 0.2 | 0.1 | 0.1 |
| intMCs | 1 | 1 | 1 | 1 | 1 | 1 | 1 | 0.2 | 0.2 | 0.2 |
| M-MDSC | 1 | 1 | 1 | 1 | 1 | 1 | 1 | 0.2 | 0.2 | 0.2 |
| mDCs | 0.9 | 0.8 | 0.8 | 1 | 0.8 | 1 | 0.8 | 0.2 | 0.1 | 0.1 |
| ncMCs | 1 | 1 | 1 | 1 | 1 | 1 | 1 | 0.2 | 0.2 | 0.2 |
| pDCs | 0 | 0 | 0 | 0 | 0 | 0 | 0 | 0 | 0 | 0 |
| $Tbet^{+}CD4^{+}Tcells_{mem}$ | 0 | 0 | 0 | 0 | 0 | 0 | 0 | 0 | 0 | 0 |
| $Tbet^{+}CD8^{+}Tcells_{mem}$ | 0 | 0 | 0 | 0 | 0 | 0 | 0 | 0 | 0 | 0 |
| $Tbet^{+}CD8^{+}Tcells_{naive}$ | 0 | 0 | 0 | 0 | 0 | 0 | 0 | 0 | 0 | 0 |
| $TCR\gamma\delta^{+}$Tcells | 0.2 | 0.2 | 0.2 | 0.2 | 0.2 | 0.2 | 0.2 | 0 | 0 | 0 |
| Tregs | 0.2 | 0.2 | 0.2 | 0.2 | 0.2 | 0.2 | 0.2 | 0 | 0 | 0 |

**Table 2. Prior knowledge matrix for prediction of Chronic Periodontitis.**

| IFNa | pCREB | pERK1/2 | IkB | pMK2 | pNFkB | pP38 | pS6 | pSTAT1 | pSTAT3 | pSTAT5 | pSTAT6 |
|---|---|---|---|---|---|---|---|---|---|---|---|
| Granulocytes | 0.4 | 0.4 | 0.4 | 0.4 | 0.4 | 0.4 | 0.2 | 1 | 0.8 | 0.7 | 0.5 |
| cMCs | 0.4 | 0.4 | 0.4 | 0.4 | 0.4 | 0.4 | 0.2 | 1 | 0.8 | 0.7 | 0.7 |
| ncMCs | 0.4 | 0.4 | 0.4 | 0.4 | 0.4 | 0.4 | 0.2 | 1 | 0.8 | 0.7 | 0.7 |
| intMCs | 0.4 | 0.4 | 0.4 | 0.4 | 0.4 | 0.4 | 0.2 | 1 | 0.8 | 0.7 | 0.7 |
| M-MDSC | 0.4 | 0.4 | 0.4 | 0.4 | 0.4 | 0.4 | 0.2 | 1 | 0.8 | 0.7 | 0.7 |
| mDCs | 0.4 | 0.4 | 0.4 | 0.4 | 0.4 | 0.4 | 0.2 | 1 | 0.8 | 0.7 | 0.7 |
| pDCs | 0.4 | 0.4 | 0.4 | 0.4 | 0.4 | 0.4 | 0.2 | 1 | 0.8 | 0.7 | 0.7 |
| $CD7^{+}$NKcells | 0.4 | 0.4 | 0.4 | 0.4 | 0.4 | 0.4 | 0.2 | 1 | 0.8 | 0.7 | 0.6 |
| $CD16^{+}CD56^{-}$NKcells | 0.4 | 0.4 | 0.4 | 0.4 | 0.4 | 0.4 | 0.2 | 1 | 0.8 | 0.7 | 0.6 |
| $CD56^{+}CD16^{-}$NKcells | 0.4 | 0.4 | 0.4 | 0.4 | 0.4 | 0.4 | 0.2 | 1 | 0.8 | 0.7 | 0.6 |

| | | | | | | | | | | | |
|---|---|---|---|---|---|---|---|---|---|---|---|
| CD4$^+$Tcells$_{mem}$ | 0.4 | 0.4 | 0.4 | 0.4 | 0.4 | 0.4 | 0.2 | 1 | 0.8 | 0.7 | 0.6 |
| CD4$^+$Tcells$_{naive}$ | 0.4 | 0.4 | 0.4 | 0.4 | 0.4 | 0.4 | 0.2 | 1 | 0.8 | 0.7 | 0.7 |
| Tbet$^+$CD4$^+$Tcells$_{mem}$ | 0.4 | 0.4 | 0.4 | 0.4 | 0.4 | 0.4 | 0.2 | 1 | 0.8 | 0.7 | 0.7 |
| Tbet$^+$CD4$^+$CD45RA$^+$Tcells | 0.4 | 0.4 | 0.4 | 0.4 | 0.4 | 0.4 | 0.2 | 1 | 0.8 | 0.7 | 0.7 |
| Tregs | 0.4 | 0.4 | 0.4 | 0.4 | 0.4 | 0.4 | 0.2 | 1 | 0.8 | 0.7 | 0.7 |
| CD8$^+$Tcells$_{mem}$ | 0.4 | 0.4 | 0.4 | 0.4 | 0.4 | 0.4 | 0.2 | 1 | 0.8 | 0.7 | 0.6 |
| CD8$^+$Tcells$_{naive}$ | 0.4 | 0.4 | 0.4 | 0.4 | 0.4 | 0.4 | 0.2 | 1 | 0.8 | 0.7 | 0.6 |
| Bcells | 0.4 | 0.4 | 0.4 | 0.4 | 0.4 | 0.4 | 0.2 | 1 | 0.8 | 0.7 | 0.7 |

| IL Cocktail | pCREB | pERK1/2 | IkB | pMK2 | pNFkB | pP38 | pS6 | pSTAT1 | pSTAT3 | pSTAT5 | pSTAT6 |
|---|---|---|---|---|---|---|---|---|---|---|---|
| Granulocytes | 0.4 | 0.4 | 0.4 | 0.4 | 0.4 | 0.4 | 0.2 | 1 | 0.8 | 0.7 | 0.5 |
| cMCs | 0.4 | 0.4 | 0.4 | 0.4 | 0.4 | 0.4 | 0.2 | 1 | 0.8 | 0.7 | 0.7 |
| ncMCs | 0.4 | 0.4 | 0.4 | 0.4 | 0.4 | 0.4 | 0.2 | 1 | 0.8 | 0.7 | 0.7 |
| intMCs | 0.4 | 0.4 | 0.4 | 0.4 | 0.4 | 0.4 | 0.2 | 1 | 0.8 | 0.7 | 0.7 |
| M-MDSC | 0.4 | 0.4 | 0.4 | 0.4 | 0.4 | 0.4 | 0.2 | 1 | 0.8 | 0.7 | 0.7 |
| mDCs | 0.4 | 0.4 | 0.4 | 0.4 | 0.4 | 0.4 | 0.2 | 1 | 0.8 | 0.7 | 0.7 |
| pDCs | 0.4 | 0.4 | 0.4 | 0.4 | 0.4 | 0.4 | 0.2 | 1 | 0.8 | 0.7 | 0.7 |
| CD7$^+$NKcells | 0.4 | 0.4 | 0.4 | 0.4 | 0.4 | 0.4 | 0.2 | 1 | 0.8 | 0.7 | 0.6 |
| CD16$^+$CD56$^-$NKcells | 0.4 | 0.4 | 0.4 | 0.4 | 0.4 | 0.4 | 0.2 | 1 | 0.8 | 0.7 | 0.6 |
| CD56$^+$CD16$^-$NKcells | 0.4 | 0.4 | 0.4 | 0.4 | 0.4 | 0.4 | 0.2 | 1 | 0.8 | 0.7 | 0.6 |
| CD4$^+$Tcells$_{mem}$ | 0.4 | 0.4 | 0.4 | 0.4 | 0.4 | 0.4 | 0.2 | 1 | 0.8 | 0.7 | 0.6 |
| CD4$^+$Tcells$_{naive}$ | 0.4 | 0.4 | 0.4 | 0.4 | 0.4 | 0.4 | 0.2 | 1 | 0.8 | 0.7 | 0.7 |
| Tbet$^+$CD4$^+$Tcells$_{mem}$ | 0.4 | 0.4 | 0.4 | 0.4 | 0.4 | 0.4 | 0.2 | 1 | 0.8 | 0.7 | 0.7 |
| Tbet$^+$CD4$^+$CD45RA$^+$Tcells | 0.4 | 0.4 | 0.4 | 0.4 | 0.4 | 0.4 | 0.2 | 1 | 0.8 | 0.7 | 0.7 |
| Tregs | 0.4 | 0.4 | 0.4 | 0.4 | 0.4 | 0.4 | 0.2 | 1 | 0.8 | 0.7 | 0.7 |
| CD8$^+$Tcells$_{mem}$ | 0.4 | 0.4 | 0.4 | 0.4 | 0.4 | 0.4 | 0.2 | 1 | 0.8 | 0.7 | 0.6 |
| CD8$^+$Tcells$_{naive}$ | 0.4 | 0.4 | 0.4 | 0.4 | 0.4 | 0.4 | 0.2 | 1 | 0.8 | 0.7 | 0.6 |
| Bcells | 0.4 | 0.4 | 0.4 | 0.4 | 0.4 | 0.4 | 0.2 | 1 | 0.8 | 0.7 | 0.7 |

| P. gingivalis | pCREB | pERK1/2 | IkB | pMK2 | pNFkB | pP38 | pS6 | pSTAT1 | pSTAT3 | pSTAT5 | pSTAT6 |
|---|---|---|---|---|---|---|---|---|---|---|---|
| Granulocytes | 0.7 | 0.7 | 0.7 | 0.7 | 0.7 | 0.7 | 0.7 | 0.2 | 0.1 | 0.1 | 0.1 |
| cMCs | 1 | 1 | 1 | 1 | 1 | 1 | 1 | 0.2 | 0.2 | 0.2 | 0.2 |
| ncMCs | 1 | 1 | 1 | 1 | 1 | 1 | 1 | 0.2 | 0.2 | 0.2 | 0.2 |
| intMCs | 1 | 1 | 1 | 1 | 1 | 1 | 1 | 0.2 | 0.2 | 0.2 | 0.2 |
| M-MDSC | 0.9 | 0.8 | 0.8 | 0.9 | 0.8 | 1 | 0.8 | 0.2 | 0.2 | 0.2 | 0.2 |
| mDCs | 0.9 | 0.8 | 0.8 | 1 | 0.8 | 1 | 0.8 | 0.2 | 0.1 | 0.1 | 0.1 |
| pDCs | 0.3 | 0.3 | 0.3 | 0.3 | 0.3 | 0.3 | 0.3 | 0 | 0 | 0 | 0 |
| CD7$^+$NKcells | 0.7 | 0.7 | 0.7 | 0.7 | 0.7 | 0.7 | 0.7 | 0 | 0 | 0 | 0 |
| CD16$^+$CD56$^-$NKcells | 0.7 | 0.7 | 0.7 | 0.7 | 0.7 | 0.7 | 0.7 | 0 | 0 | 0 | 0 |

| | | | | | | | | | | | |
|---|---|---|---|---|---|---|---|---|---|---|---|
| CD56$^+$CD16$^-$NKcells | 0.7 | 0.7 | 0.7 | 0.7 | 0.7 | 0.7 | 0.7 | 0 | 0 | 0 | 0 |
| CD4$^+$Tcells$_{mem}$ | 0 | 0 | 0 | 0 | 0 | 0 | 0 | 0 | 0 | 0 | 0 |
| CD4$^+$Tcells$_{naive}$ | 0 | 0 | 0 | 0 | 0 | 0 | 0 | 0 | 0 | 0 | 0 |
| Tbet$^+$CD4$^+$Tcells$_{mem}$ | 0 | 0 | 0 | 0 | 0 | 0 | 0 | 0 | 0 | 0 | 0 |
| Tbet$^+$CD4$^+$CD45RA$^+$Tcells | 0 | 0 | 0 | 0 | 0 | 0 | 0 | 0 | 0 | 0 | 0 |
| Tregs | 0.2 | 0.2 | 0.2 | 0.2 | 0.2 | 0.2 | 0.2 | 0 | 0 | 0 | 0 |
| CD8$^+$Tcells$_{mem}$ | 0 | 0 | 0 | 0 | 0 | 0 | 0 | 0 | 0 | 0 | 0 |
| CD8$^+$Tcells$_{naive}$ | 0 | 0 | 0 | 0 | 0 | 0 | 0 | 0 | 0 | 0 | 0 |
| Bcells | 0.2 | 0.2 | 0.2 | 0.2 | 0.2 | 0.2 | 0.2 | 0 | 0 | 0 | 0 |

| TNFa | pCREB | pERK1/2 | IkB | pMK2 | pNFkB | pP38 | pS6 | pSTAT1 | pSTAT3 | pSTAT5 | pSTAT6 |
|---|---|---|---|---|---|---|---|---|---|---|---|
| Granulocytes | 0.8 | 0.7 | 0.8 | 0.8 | 0.8 | 0.8 | 0.7 | 0.1 | 0.2 | 0.2 | 0.2 |
| cMCs | 0.9 | 0.8 | 1 | 0.8 | 1 | 0.8 | 0.8 | 0.1 | 0.2 | 0.2 | 0.2 |
| ncMCs | 0.9 | 0.8 | 1 | 0.8 | 1 | 0.8 | 0.8 | 0.1 | 0.2 | 0.2 | 0.2 |
| intMCs | 0.9 | 0.8 | 1 | 0.8 | 1 | 0.9 | 0.8 | 0.1 | 0.2 | 0.2 | 0.2 |
| M-MDSC | 0.8 | 0.8 | 1 | 0.8 | 1 | 0.8 | 0.8 | 0.1 | 0.2 | 0.2 | 0.2 |
| mDCs | 0.9 | 0.8 | 1 | 0.8 | 1 | 0.8 | 0.8 | 0.1 | 0.2 | 0.2 | 0.2 |
| pDCs | 0.8 | 0.8 | 1 | 0.8 | 1 | 0.8 | 0.8 | 0.1 | 0.2 | 0.2 | 0.2 |
| CD7$^+$NKcells | 0.8 | 0.8 | 1 | 0.8 | 1 | 0.8 | 0.8 | 0.1 | 0.2 | 0.2 | 0.2 |
| CD16$^+$CD56$^-$NKcells | 0.8 | 0.8 | 1 | 0.8 | 1 | 0.8 | 0.8 | 0.1 | 0.2 | 0.2 | 0.2 |
| CD56$^+$CD16$^-$NKcells | 0.8 | 0.8 | 1 | 0.8 | 1 | 0.8 | 0.8 | 0.1 | 0.2 | 0.2 | 0.2 |
| CD4$^+$Tcells$_{mem}$ | 0.7 | 0.7 | 0.7 | 0.7 | 0.7 | 0.7 | 0.7 | 0.1 | 0.2 | 0.2 | 0.2 |
| CD4$^+$Tcells$_{naive}$ | 0.7 | 0.7 | 0.7 | 0.7 | 0.7 | 0.7 | 0.7 | 0.1 | 0.2 | 0.2 | 0.2 |
| Tbet$^+$CD4$^+$Tcells$_{mem}$ | 0.7 | 0.7 | 0.7 | 0.7 | 0.7 | 0.7 | 0.7 | 0.1 | 0.2 | 0.2 | 0.2 |
| Tbet$^+$CD4$^+$CD45RA$^+$Tcells | 0.7 | 0.7 | 0.7 | 0.7 | 0.7 | 0.7 | 0.7 | 0.1 | 0.2 | 0.2 | 0.2 |
| Tregs | 0.7 | 0.7 | 0.7 | 0.7 | 0.7 | 0.7 | 0.7 | 0.1 | 0.2 | 0.2 | 0.2 |
| CD8$^+$Tcells$_{mem}$ | 0.7 | 0.7 | 0.7 | 0.7 | 0.7 | 0.7 | 0.7 | 0.1 | 0.2 | 0.2 | 0.2 |
| CD8$^+$Tcells$_{naive}$ | 0.7 | 0.7 | 0.7 | 0.7 | 0.7 | 0.7 | 0.7 | 0.1 | 0.2 | 0.2 | 0.2 |
| Bcells | 0.7 | 0.7 | 0.7 | 0.7 | 0.7 | 0.7 | 0.7 | 0.1 | 0.2 | 0.2 | 0.2 |

**Table 3. Panel used for the mass cytometry assay of chronic periodontitis patient whole blood samples.**

| Antibody | Manufacturer | Symbol | Atomic Mass | Clone | Comment |
|---|---|---|---|---|---|
| Barcode 1 | Trace Sciences | Pd | 102 | | Barcode |
| Barcode 2 | Trace Sciences | Pd | 104 | | Barcode |

| | | | | | |
|---|---|---|---|---|---|
| Barcode 3 | Trace Sciences | Pd | 105 | | Barcode |
| Barcode 4 | Trace Sciences | Pd | 106 | | Barcode |
| Barcode 5 | Trace Sciences | Pd | 108 | | Barcode |
| Barcode 6 | Trace Sciences | Pd | 110 | | Barcode |
| CD235ab | Biolegend | In | 113 | HIR2 | Phenotype |
| CD61 | BD | In | 113 | VI-PL2 | Phenotype |
| CD45 | Biolegend | In | 115 | HI30 | Phenotype |
| CD66 | BD | La | 139 | CD66a-B1.1 | Phenotype |
| CD7 | BD | Pr | 141 | M-T701 | Phenotype |
| CD19 | Biolegend | Nd | 142 | HIB19 | Phenotype |
| CD45RA | Biolegend | Nd | 143 | HI100 | Phenotype |
| CD11b | Fluidigm | Nd | 144 | ICRF44 | Phenotype |
| CD4 | Fluidigm | Nd | 145 | RPA-T4 | Phenotype |
| CD8a | Fluidigm | Nd | 146 | RPA-T8 | Phenotype |
| CD11c | Fluidigm | Sm | 147 | Bu15 | Phenotype |
| CD123 | Biolegend | Nd | 148 | 6H6 | Phenotype |
| pCREB | Cell Signaling Technology | Sm | 149 | 87G3 | Phenotype |
| pSTAT5 | Fluidigm | Nd | 150 | 47 | Function |
| pp38 | BD | Eu | 151 | 36/p38 | Function |
| TCRγδ | Fluidigm | Sm | 152 | 11F2 | Phenotype |
| pSTAT1 | Fluidigm | Eu | 153 | 58D6 | Function |
| pSTAT3 | Cell Signaling Technology | Sm | 154 | M9C6 | Function |
| pS6 | Cell Signaling Technology | Gd | 155 | D57.2.2E | Function |
| CD24 | Biolegend | Gd | 156 | ML5 | Phenotype |
| CD38 | Biolegend | Gd | 157 | HIT2 | Phenotype |
| CD33 | Fluidigm | Gd | 158 | WM53 | Phenotype |
| pMAPKAPK2 | Fluidigm | Tb | 159 | 27B7 | Function |
| Tbet | Fluidigm | Gd | 160 | 4B10 | Function |

| | | | | | |
|---|---|---|---|---|---|
| cPARP | BD | Dy | 161 | F21-852 | Function |
| FoxP3 | Fluidigm | Dy | 162 | PCH101 | Phenotype |
| IκB | Fluidigm | Dy | 164 | L35A5 | Function |
| CD16 | Fluidigm | Ho | 165 | 3G8 | Phenotype |
| pNF-κB | Fluidigm | Er | 166 | K10-895.12.50 | Function |
| pERK1/2 | Fluidigm | Er | 167 | D13.14.4E | Function |
| pSTAT6 | Fluidigm | Er | 168 | 18 | Function |
| CD25 | Biolegend | Tm | 169 | M-A251 | Phenotype |
| CD3 | Fluidigm | Er | 170 | UCHT1 | Phenotype |
| CD27 | BD | Yb | 171 | M-T271 | Phenotype |
| CCR2 | Biolegend | Yb | 173 | K036C2 | Phenotype |
| HLA-DR | Fluidigm | Yb | 174 | L243 | Phenotype |
| CD56 | BD | Yb | 176 | NCAM16.2 | Phenotype |
| DNA1 | Fluidigm | Ir | 191 | | DNA |
| DNA2 | Fluidigm | Ir | 192 | | DNA |

**Table 4. Panel used for the mass cytometry assay of pregnant women whole blood samples.**

| Antibody | Manufacturer | Symbol | Atomic Mass | Clone | Comment |
|---|---|---|---|---|---|
| Barcode 1 | Trace Sciences | Pd | 102 | | Barcode |
| Barcode 2 | Trace Sciences | Pd | 104 | | Barcode |
| Barcode 3 | Trace Sciences | Pd | 105 | | Barcode |
| Barcode 4 | Trace Sciences | Pd | 106 | | Barcode |
| Barcode 5 | Trace Sciences | Pd | 108 | | Barcode |
| Barcode 6 | Trace Sciences | Pd | 110 | | Barcode |
| CD235ab | Biolegend | In | 113 | HIR2 | Phenotype |
| CD61 | BD | In | 113 | VI-PL2 | Phenotype |
| CD45 | Biolegend | In | 115 | HI30 | Phenotype |
| CD66 | BD | La | 139 | CD66a-B1.1 | Phenotype |

| | | | | | |
|---|---|---|---|---|---|
| CD7 | BD | Pr | 141 | M-T701 | Phenotype |
| CD19 | Fluidigm | Nd | 142 | HIB19 | Phenotype |
| CD45RA | Fluidigm | Nd | 143 | HI100 | Phenotype |
| CD11b | Fluidigm | Nd | 144 | ICRF44 | Phenotype |
| CD4 | Fluidigm | Nd | 145 | RPA-T4 | Phenotype |
| CD8a | Fluidigm | Nd | 146 | RPA-T8 | Phenotype |
| CD11c | Fluidigm | Sm | 147 | Bu15 | Phenotype |
| CD123 | Biolegend | Nd | 148 | 6H6 | Phenotype |
| pCREB | Cell Signaling Technology | Sm | 149 | 87G3 | Phenotype |
| pSTAT5 | Fluidigm | Nd | 150 | 47 | Function |
| pp38 | CST | Eu | 151 | 36/p38/pT18 | Function |
| TCRγδ | Fluidigm | Sm | 152 | 11F2 | Phenotype |
| pSTAT1 | Fluidigm | Eu | 153 | 58D6 | Function |
| pSTAT3 | BD | Sm | 154 | 4/P pY705 | Function |
| pS6 | Cell Signaling Technology | Gd | 155 | D57.2.2E | Function |
| CD33 | Fluidigm | Gd | 158 | WM53 | Phenotype |
| pMAPKAPK2 | Fluidigm | Tb | 159 | 27B7 | Function |
| Tbet | Fluidigm | Gd | 160 | 4B10 | Phenotype |
| FoxP3 | Fluidigm | Dy | 162 | PCH101 | Phenotype |
| IκB | Fluidigm | Dy | 164 | L35A5 | Function |
| CD16 | Fluidigm | Ho | 165 | 3G8 | Phenotype |
| pNF-κB | Fluidigm | Er | 166 | K10-895.12.50 | Function |
| pERK1/2 | CST | Er | 167 | D13.14.4E | Function |
| CD25 | Biolegend | Tm | 169 | M-A251 | Phenotype |
| CD3 | Fluidigm | Er | 170 | UCHT1 | Phenotype |
| CD15 | Fluidigm | Yb | 172 | W6D3 | Phenotype |
| HLA-DR | Fluidigm | Yb | 174 | L243 | Phenotype |
| CD14 | Fluidigm | Yb | 175 | M52E | Phenotype |
| CD56 | Fluidigm | Yb | 176 | NCAM16.2 | Phenotype |

| | | | | | |
|---|---|---|---|---|---|
| DNA1 | Fluidigm | Ir | 191 | | DNA |
| DNA2 | Fluidigm | Ir | 192 | | DNA |
